# Late endosome transport by RILP-RAB7A promotes dendrite arborization

**DOI:** 10.1101/2025.09.03.673267

**Authors:** Chan Choo Yap, Laura Digilio, Lloyd P. McMahon, Ryan J. Mulligan, Isabelle F. Witteveen, Bettina Winckler

## Abstract

Directional dendritic transport of late endosomes retrogradely towards the soma is required for fusion with lysosomes and for degradation in the soma. Both dendritic motility of late endosomes and somatic degradation require RAB7A. Similarly, interference with dynein function reduces motility of late endosomes and results in degradative failure. Blocking dynein function also impairs normal dendrite growth, suggesting that motility of late endosomes and/or lysosomes might be required for dendrite growth. RAB7A and dynein are mechanistically linked via RILP which is a dynein-interacting RAB7A effector. RILP also binds the late endosome-lysosome fusion tether HOPS. In non-neuronal cells, downregulation of RILP leads to impaired degradation due to deficiencies in late endosome transport and fusion defects with lysosomes. In this work, we express a separation-of-function mutant of RAB7A (RAB7A-L8A) incapable of RILP binding. Based on the results in non-neuronal cells, we hypothesized that both endosome motility and degradation in neurons depended on RILP. Our data in cultured rat and mouse hippocampal neurons of both sexes suggest that endogenous RILP is a functional RAB7A-dependent dynein adaptor for late endosome motility in dendrites. Interestingly, it also promotes endosome carrier formation. As a consequence of late endosome transport inhibition, degradative cargos are not cleared normally from dendrites in RAB7A-L8A. Surprisingly, lysosomal fusion and somatic degradation do not require RAB7A-RILP interactions. Despite the normal degradation, dendrite arborization is impaired in RAB7A-L8A expressing neurons, demonstrating that dendrite morphology defects are separable from degradation blockade. This indicates that normal dendrite growth/maintenance is dependent on sustained RAB7A/RILP-dependent LE transport.

**Significance Statement:** Dendrite growth requires membrane trafficking, but the roles of individual compartments and regulators are not well established. Stunted dendrite growth is often associated with endolysosomal traffic jams and degradation block. In contrast, our work reveals a requirement for transport of late endosomes to support dendrite growth independently of late endosomes as carriers of degradative cargos to the lysosome for degradation.

## Introduction

Vesicular transport along dendrites is well documented (Yap and Winckler, 2022). Yet, unlike for directional axonal transport (for which the purpose is well established), the regulation of directional dendritic transport is less well understood. Directional dendritic transport is also conceptually complicated since microtubule polarity is close to 50:50 plus-end out vs minus-end out (Kapitein and Hoogenraad, 2011; Yau et al., 2016; Masucci et al., 2021; Baas and Lin, 2011), and kinesins and dynein both have been implicated in anterograde and retrograde transport (Ayloo et al., 2017; Kapitein et al., 2010; Tas et al., 2017). One process that does require directional vesicular transport in dendrites is the bulk degradation of membrane cargos (Yap et al., 2018). Because the vast majority of lysosomes reside in the soma and in proximal dendrites, and far fewer lysosomes are found further out along dendrites (Yap et al., 2022c), degradative dendritic cargos need to transport retrogradely to the soma for fusion with lysosomes. In addition, growth and maintenance of dendrites require organelle transport. For instance, dendritic receptors that regulate dendrite morphogenesis (such as growth factor and guidance cue receptors) require anterograde transport of TGN-derived organelles and distal exocytosis (Santana and Marzolo, 2017; Kennedy and Hanus, 2019; Radler et al., 2020; Wang et al., 2012; Horton et al., 2005; Bowen et al., 2017). Furthermore, regulation of dendritic receptor levels relies on endocytosis and subsequent trafficking of endosomes (Moya-Alvarado et al., 2022; Ayloo et al., 2017; Yap et al., 2022a; Tang et al., 2020; González-Gutiérrez et al., 2020; Kanamori et al., 2015).

Proteostasis in all cells is regulated by the endo-lysosomal system which degrades proteins entering cells via endocytosis (Klumperman and Raposo, 2014; Winckler et al., 2018; Britt et al., 2016) and intracellular proteins via autophagy (Sidibe et al., 2022; Vargas et al., 2023; Boecker and Holzbaur, 2019). In addition, signaling duration of a multitude of endocytosed receptors (for example Trk receptors) is set by the rate with which they leave LEs (where they signal) and reach lysosomes (where they degrade) (Winckler and Yap, 2011; Barford et al., 2017; Suo et al., 2014). Not surprisingly, genes linked to endosomal and lysosomal pathways are frequently associated with diseases of the nervous system (Nixon, 2024; Ferguson, 2019). Endocytosed cargos reach the lysosomes via multiple endosomal intermediates, specifically early endosomes (EEs) which mature to RAB7A-positive late endosomes (LEs) which fuse with lysosomes (Lys) (Klumperman and Raposo, 2014). Endosomal-lysosomal pathways are controlled by a large number of proteins, including the small GTPase RAB7A, to regulate flux through the endosomal system and maintain protein homeostasis and proper signaling levels emanating from EEs and LEs (Yap et al., 2022c; Klumperman and Raposo, 2014). Of note, RAB7A mutations are linked to Charcot-Marie-Tooth disease 2B and have been implicated in dysregulating neurotrophin signaling (Mulligan and Winckler, 2023; Guerra and Bucci, 2016). Importantly, RAB7A function depends on a large number of effector proteins which mediate RAB7A function in endosome maturation, motility, and fusion (Mulligan and Winckler, 2023; Wang et al., 2011). The precise roles of many RAB7A effectors is still being uncovered.

RILP is a multifunctional RAB7A effector whose interactors include dynein and fusion machinery (HOPS) among others (Vallee et al., 2021; Wang et al., 2011; De Luca and Bucci, 2014; Jongsma et al., 2024; Rocha et al., 2009; Progida et al., 2006; Lin et al., 2014; Cantalupo et al., 2001). In past work, we identified dynein as a retrograde motor for dendritic LEs (Yap et al., 2022a). We showed that overexpression of RILP clustered all RAB7A-positive compartments in the soma. Furthermore, mild overexpression of RILP led to increased net retrograde motility of LEs whereas inhibition of dynein reduced motility. In support of the notion that degradation of dendritic receptors requires retrograde transport, RAB7A and dynein function were both required for normal degradation of the short-lived dendritic receptors NSG1/2. We now test whether endogenous RILP is the dendritic dynein adaptor for LEs.

## Results

### Localization of myc-RILP in dendritic late endosomes and somatic lysosomes

We first sought to determine which endosomes in soma and dendrites were associated with RILP. We tried several commercially available anti-RILP antibodies, but none resulted in reliable staining. Two of the most commonly used anti-RILP antibodies only recognized human, but not mouse or rat RILP (Yap et al., 2023, 2022b). In addition, transcriptome data bases show that RILP mRNA is expressed at very low levels (brainrnaseq.org), making detection of endogenous RILP challenging. We thus switched to a short overexpression paradigm to visualize RILP localization since high RILP overexpression changed LE and lysosome positioning (Yap et al., 2022a). Myc-RILP expressed together with GFP-RAB7A showed largely co-localized patterns in the soma and dendrites (Fig. 1A), consistent with RILP being a RAB7A effector.

**Figure 1:**
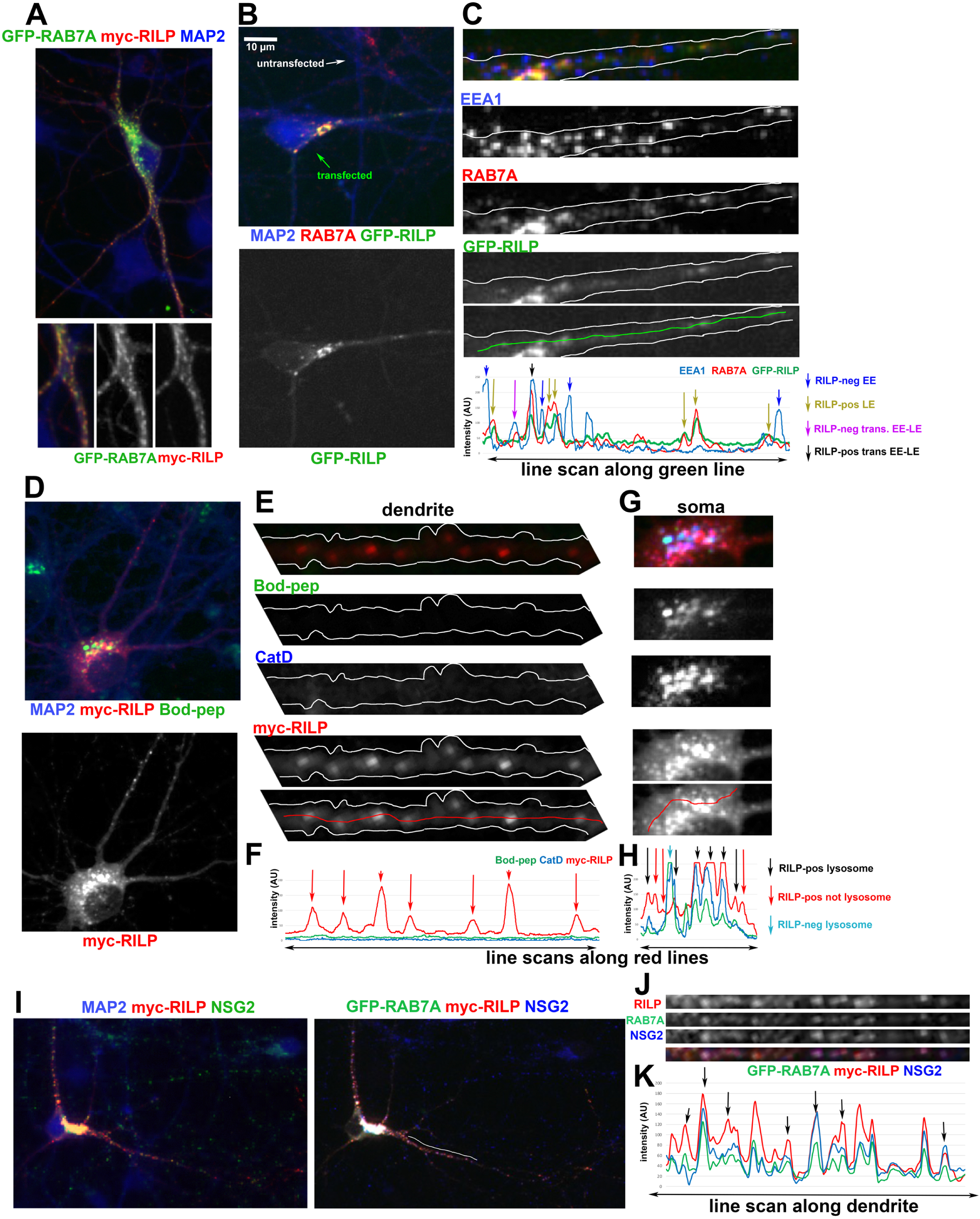
Localization of tagged RILP in dendritic late endosomes and somatic lysosomes. (A-C) Tagged RILP (expressed for 8 hours) localizes with late endosomes, detected by GFP-RAB7A (A) or endogenous RAB7A (B,C). Individual channels along a major dendrite are shown in (C) with the corresponding line scans. Arrows indicate RILP-negative early endosomes EE (blue), RILP-positive late endosomes LE (yellow) and a mixture of RILP-positive (black) and RILP-negative (magenta) transitioning EE-LEs (tEE-LE).sarala(D-H) Briefly expressed tagged RILP (red) localizes with lysosomes, detected by Bodipy-pepstatin (green; D) or endogenous cathepsin D, CatD (blue) (E,G). Individual channels along a major dendrite (E) and the soma (G) with the corresponding line scans (F,H) are shown. Along dendrites (F), RILP-positive compartments (red arrows) do not contain lysosomal markers. In the soma (H), arrows indicate lysosomes that either do (black arrows) or do not (blue arrows) contain RILP.sarala(I-K) Briefly expressed tagged RILP (red) localizes with the degradative cargo NSG2 (blue) and GFP-RAB7A (green). The whole cell is shown in (I), single channels of one dendrite are shown in (J) and the corresponding line scan is shown in (K). Black arrows point at co-localized compartments along dendrites. MAP2 (blue) indicates dendrites in (A,B, D, I). All images are of DIV10 rat hippocampal neurons.

We then determined if RILP is already associated with transitioning EE-LEs (tEE-LE: double-positive for EEA1 and RAB7A). GFP-RILP expressed for 8 hours and then co-stained against endogenous EEA1 (to demarcate EEs) and endogenous RAB7A (to demarcate LEs; Fig. 1B) did not co-localize highly with EEA1 alone but localized well to RAB7A-positive LEs (Fig. 1C). It also localized to some but not all tEE-LEs (Fig. 1C; black arrows). These transitioning compartments constitute about 25% of RAB7A compartments in dendrites and are a maturational intermediate between EEs and LEs (Yap et al., 2018). RILP thus starts to associate with tEE-LEs before EEA1 dissociates.

In order to determine if RILP remained associated with degradative lysosomes, we co-stained neurons transfected for 8 hours with myc-RILP against the lysosomal protease Cathepsin D (CatD) after live incubation with bodipy-pepstatin (Bod-pep) which marks acidified, degradatively active lysosomes (Fig. 1D-H). Myc-RILP containing compartments could be found along dendrites, but these compartments did not contain detectable CatD or Bod-pep (Fig. 1E,F) and thus correspond to LEs and tEE-LEs. In contrast, somatic compartments containing myc-RILP often corresponded to degradative lysosomes (Fig. 1G,H). We note that we also observed a subset of somatic degradative lysosomes that did not contain myc-RILP (Fig. 1H). In order to further confirm that most of the dendritic myc-RILP containing compartments in dendrites were LEs, we co-stained myc-RILP expressing neurons against the short-lived dendritic receptor NSG2 (Fig. 1I) which is overwhelmingly found in LEs as it travels retrogradely to somatic lysosomes (Yap et al., 2017). We find very high co-localization of myc-RILP with GFP-RAB7A and NSG2 along dendrites (Fig.1 J,K). We thus find that RILP localizes to dendritic and somatic tEE-LEs, LEs, and on a subset of somatic lysosomes.

### Characterization of RILP-binding mutants RAB7A-L8A and RAB7A-F45A

We initially used knockdown approaches against RILP in order to ask whether loss of RILP affects motility of LEs and/or degradation but encountered pervasive off-target effects in neurons (Yap et al., 2023). As an alternative approach, we now generated separation-of-function mutants of RAB7A that no longer bind to RILP (RAB7A-L8A and RAB7A-F45A (Wu et al., 2005; Lin et al., 2014)). We confirmed that neither of them co-IPed with myc-RILP (SFig. 1A), similar to the dominant-negative RAB7A-T22N (“RAB7A-DN”) which lacks binding to all its effectors. WT GFP-RAB7A, as expected, co-immunoprecipitated with myc-RILP (SFig. 1A). Quantification of the co-IP (SFig. 1B) showed that both mutants retained <5% of the RILP-binding capacity of WT RAB7A.

In addition to co-IPs, we transfected neuronal cultures with WT RAB7A or its mutants together with myc-RILP to determine if RILP was still recruited to endosomes. siRAB7 was co-transfected to reduce the levels of endogenous RAB7A. Consistent with previous results (Yap et al., 2022a), overexpression of myc-RILP together with WT GFP-RAB7A led to strong co-labeling of endosomes (SFig. 1C). In contrast, neither Em-RAB7A-L8A nor Em-RAB7A-F45A recruited and clustered myc-RILP, leaving it diffusely cytosolic (SFig 1.D,E). This is consistent with the lack of RILP binding by RAB7-L8A and RAB7A-F45A.

### RAB7A-L8A localizes to many endosomes and lysosomes in neurons

In previous work, we observed that overexpression of RILP hyperstabilized endogenous RAB7A on endosomes leading to a >5x higher level of RAB7A on endosomes (Yap et al., 2022a). We thus hypothesized that loss of RILP binding would affect the ability of RAB7A-L8A or RAB7A-F45A to associate with endosomes. We knocked down endogenous RAB7A with siRAB7A while at the same time re-expressing WT RAB7A, RAB7A-L8A (Fig. 2) or RAB7A-F45A (SFig. 2A-D). Quantification of re-expressed RAB7A WT/L8A/F45A levels compared to endogenous RAB7A levels is shown in Fig. 2A. Similarly to WT GFP-RAB7A (Fig. 2B; green), we found clear association of Em-RAB7A-L8A with intracellular organelles (Fig. 2C; green). RAB7A-L8A can thus stably associate with intracellular organelles suggesting the possibility that it still interacts with other RAB7A effectors.

**Figure 2:**
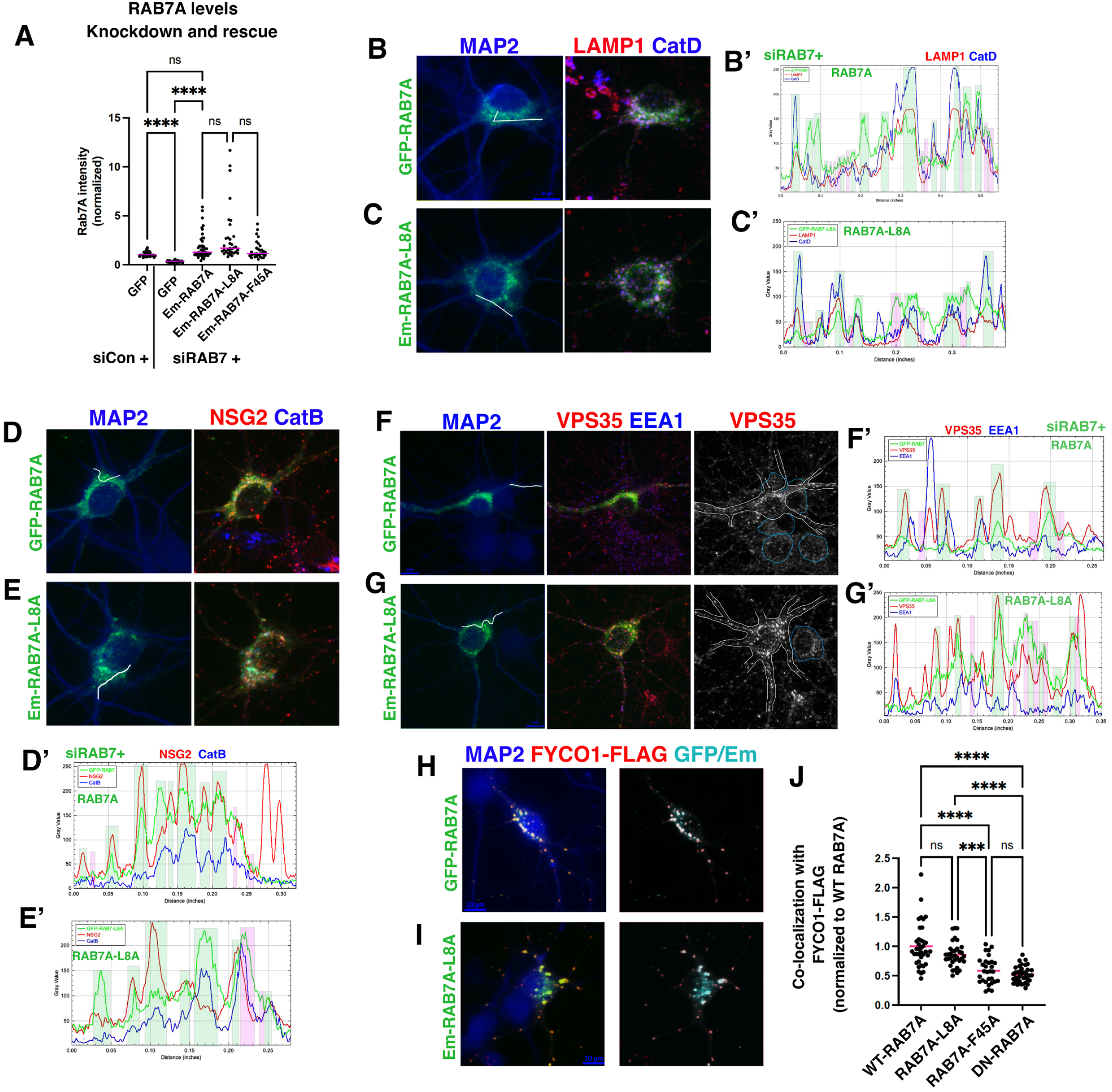
RAB7A-L8A still recruits its effectors VPS35 and FYCO1. (A) Somatic RAB7A levels were quantified in DIV10 hippocampal neurons co-transfected with si-Control + GFP or siRAB7A + GFP, WT RAB7A, RAB7A-L8A or RAB7A-F45A. The levels of rescue constructs were in the same range as the endogenous RAB7A levels. Statistics: N= 29-46 cells/condition. Kruskal-Wallis test. **** p<0.0001. Medians are shown.sarala(B-E) Localization of tagged WT RAB7A (B,D) and RAB7A-L8A (C,E) with endogenous markers of somatic endosomes (LAMP1: B,C; NSG2: D,E) and lysosomes (LAMP1: B,C; CatD: B,C; CatB: D,E) in DIV10 rat hippocampal neurons is shown together with line scans across the soma (B’-E’’): Coinciding peaks are highlighted in green whereas non-coinciding parks are pink. MAP2 marks dendrites. siRAB7 was co-transfected to minimize contribution to marker distribution from endogenous RAB7A.sarala(F,G) Localization of tagged WT RAB7A (F) and RAB7A-L8A (G) with endogenous markers of somatic early endosomes (Vps35 in red; EEA1 in blue) in DIV10 rat hippocampal neurons is shown together with line scans across the soma (F’,G’): Coinciding peaks are highlighted in green whereas non-coinciding peaks are pink. A single channel is shown for easier comparison Vps35 levels in transfected (white outline) and non-transfected (blue outline) neurons.sarala(H-J) WT tagged RAB7A (H) and RAB7A-L8A (I) (green or cyan) both localize strongly with FYCO1-FLAG (red) in transfected neurons. (J) Quantification of extent that different RAB7A proteins co-localize with FCYO1-positive compartments in DIV9 neurons transfected with FYCO1-FLAG and RAB7A constructs. The ratio of cytosolic vs FYCO1-compartment associated RAB7A fluorescence was determined and normalized to WT RAB7A. Statistics: N = 30-38 cells/condition from 2 independent experiments. Kruskal-Wallis test. Mean is indicated by pink line. *** p<0.001, **** p<0.0001.

In order to identify which compartments recruited RAB7A-L8A, we co-stained neurons co-expressing siRAB7A and GFP-RAB7A or Em-RAB7A-L8A with endogenous markers of lysosomes (CatD, CatB, LAMP1; Fig. 2B-E), LEs (LAMP1, NSG2; Fig. 2B-E), and EEs (EEA1; Fig. 2F,G). GFP-RAB7A and Em-RAB7A-L8A showed similar co-localization with a subset of EEs, LEs and Lys (see line scans Fig. 2B’-G’: co-localized peaks are highlighted by green bars, absence of co-localization highlighted by pink bars).

### RAB7A-L8A still recruits its effectors VPS35 and FYCO1

In addition, we stained against endogenous VPS35 (Fig. 2F,G) which is itself a RAB7A effector. VPS35 is part of the retromer complex, can be recruited to LEs by RAB7A and is implicated in the retrieval of several cargos to the trans-Golgi network (Priya et al., 2015; Burd and Cullen, 2014). Similar to WT RAB7A, RAB7A-L8A co-localized with a subset of VPS35-containing compartments (see line scans in Fig. 2F’,G’). Interestingly, overexpression of WT RAB7A led to increased VPS35 intensity on compartments, likely by decreased dissociation leading to hyper-stabilization. The same was observed for RAB7A-L8A (Fig. 2F,G), suggesting that RAB7A-L8A interacted similarly with VPS35 as WT RAB7A.

We then determined if RAB7A-L8A was still stabilized by another RAB7A effector, FYCO1 (Pankiv et al., 2010). FYCO1 overexpression led to bright accumulation of WT GFP-RAB7A on somatic and dendritic endosomes (Fig. 2H), again likely due to hyperstability of FYCO1-bound RAB7A. Em-RAB7A-L8A was also strongly stabilized on FYCO1-positive endosomes (Fig. 2I). Quantification of the proportion of WT RAB7A and RAB7A-L8A on FYCO1-positive compartments showed no differences (Fig. 2J), suggesting that FYCO1 recruitment is not disrupted by the L8A point mutation. We thus determined that RAB7A-L8A showed near-complete loss of RILP binding whilst maintaining associations with at least two of its other effectors, VPS35 and FYCO1. RAB7A-L8A thus acts as a “separation-of-function mutant” and can be used to ask how loss of RILP-binding by RAB7A affects RAB7A-dependent functions in neurons.

Em-RAB7A-F45A was still detectably enriched on intracellular compartments in some transfected neurons (SFig. 2A-D; green), but in many cells Em-RAB7A-F45A had an overwhelmingly cytosolic distribution with little compartment recruitment evident. In contrast to WT RAB7A and RAB7A-L8A, we did not consistently observe the hyper-recruitment of VPS35 by RAB7A-F45A in transfected cells (SFig. 2C). Furthermore, RAB7A-F45A was inefficiently accumulated on compartments with overexpressed FYCO1 (SFig. 2D) compared to WT RAB7A and RAB7A-L8A and was comparable to DN-RAB7A with respect to FYCO1 co-localization (Fig. 2J). RAB7A-F45A is thus completely deficient in RILP binding but also (partially) deficient in associating with additional RAB7A effectors.

### Loss of RILP binding by RAB7A impairs motility of dendritic late endosomes

Since RILP is a known dynein adaptor for RAB7A on LEs in fibroblasts (Johansson et al., 2007), and dynein is required for motility of LEs in dendrites (Yap et al., 2022a), we asked if RAB7-L8A could still promote normal motility of LEs in dendrites. We carried out live imaging of GFP-RAB7A WT or Em-RAB7A-L8A transfected together with siRAB7A and the LE cargo NSG1-mCh. Kymographs (Fig. 3A,B) were analyzed with KymoButler machine learning software (which we trained on RAB7A movies) to determine the net displacement of individual RAB7A-positive organelles during 400 seconds of live imaging (as in our previous work (Yap et al., 2022a). The motility of RAB7A (representing total RAB7A-positive population: tEE-LEs + LEs) and NSG1 (representing LEs as well as TGN-derived carriers) was analyzed separately. In RAB7-L8A expressing cells, the stationary pool was increased for both RAB7-L8A and NSG1 compared to WT RAB7A expressing cells (Fig. 3C; purple bar in left graph). For NSG1 carriers, there was also a small but significant reduction in retrograde (green bar: 18.5% reduced to 14.3%) but not anterograde tracks (pink bar) (Fig. 3C; right graph). This result argues that the RAB7A-RILP complex increases motility of a subset of LEs in dendrites.

**Figure 3:**
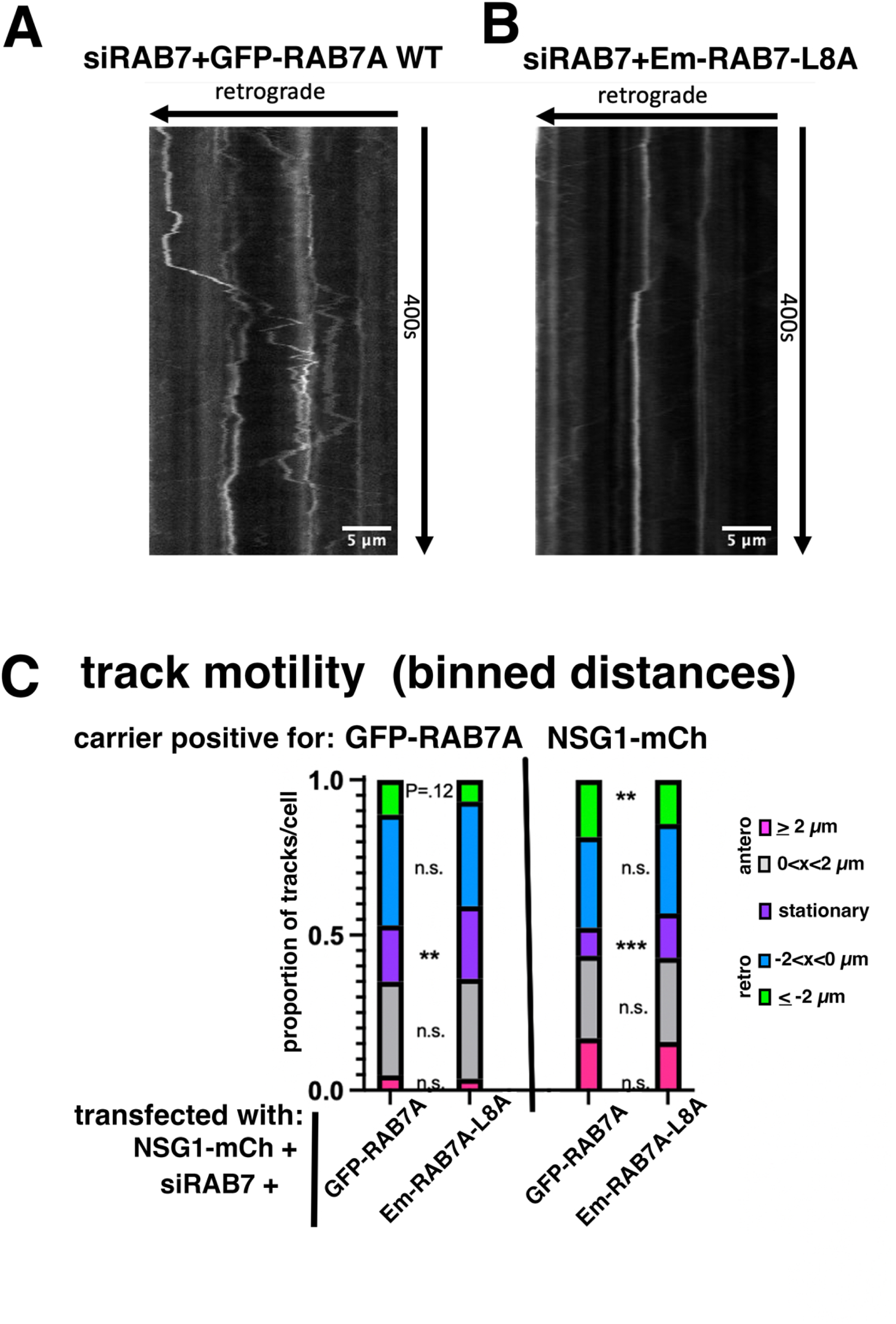
Loss of RILP binding by RAB7A impairs motility of dendritic late endosomes. (A-C) Motility of dendritic late endosomes is impaired in RAB7A-L8A expressing neurons. GFP-RAB7A (A) or Em-RAB7A-L8A (B) motility was imaged live in dendrites of DIV10 cultured rat hippocampal neurons co-transfected with WT RAB7A or RAB7A-L8A and simultaneously with the LE cargo NSG1-mCh. siRAB7 was also co-transfected to minimize contributions from endogenous RAB7A. (C) Motility of RAB7A- or NSG1-positive carriers was quantified from kymographs using Kymobutler. Net track motility was binned into 5 categories by directionality (retrograde vs anterograde vs stationary) and extent of movement (small = less than 2 µm in either direction; large = more than or equal to 2 µm in either direction), as indicated in the color-coded legend. Statistics by cell – N = 24-26 cells from 2 independent cultures representing 89-111 dendrites. Mann-Whitney or unpaired t-test, as appropriate. ** p<0.01, *** p<0.001.

### Loss of RILP binding by RAB7A decreases dendrite elaboration

Given the importance of maintaining proteostasis for neuronal health overall (Cullen et al., 2024; Malik et al., 2019; Mestres and Sung, 2017; Ivanova and Cousin, 2022; Plooster et al., 2022), we next tested whether RILP binding by RAB7A is required for dendrite growth and/or maintenance. We previously showed that inhibition of dynein led to reduction in dendrite length (Yap et al., 2022a). We thus hypothesized that loss of RILP binding in RAB7A-L8A and -F45A would similarly affect dendrite length. Inhibition of RAB7A with siRAB7 + RAB7A-DN led to greatly simplified arborization compared to siCon+WT RAB7A and siRAB7+WT RAB7A (i.e. rescue) (Fig. 4A-C) (similar to Schwenk et al., 2014). We quantified the total dendrite length in controls and in siRAB7A-downregulated neurons with or without rescue by WT RAB7A. Total dendrite length was reduced by ∼30% when RAB7A was downregulated compared to neurons transfected with a control si-plasmid (siCon) but could be completely rescued by co-expression of WT RAB7A (Fig. 4D). Neurons expressing RAB7A-DN showed reduced total dendrite length compared to GFP in the context of siCon (Fig. 4E: ∼30% reduction). RAB7A-DN further augmented the phenotype of siRAB7 alone (Fig. 4D: ∼50% reduction) demonstrating that it reduced residual RAB7A function. In contrast to RAB7A-DN, expression of either RAB7A-L8A or -F45A did not change total filament length in the context of siCon (Fig. 4E showing that they are not dominant-negative. We then compared the rescue ability of WT RAB7A (indicated by blue line in Fig. 4F) to that of RAB7A-L8A and RAB7A-F45A in the context of siRAB7 (indicated by green line in Fig. 4F). RAB7A-L8A and -F45A were not able to rescue total dendrite length (Fig. 4F; ∼20% reduction for L8A and ∼30% reduction for F45A compared to WT RAB7A).

**Figure 4:**
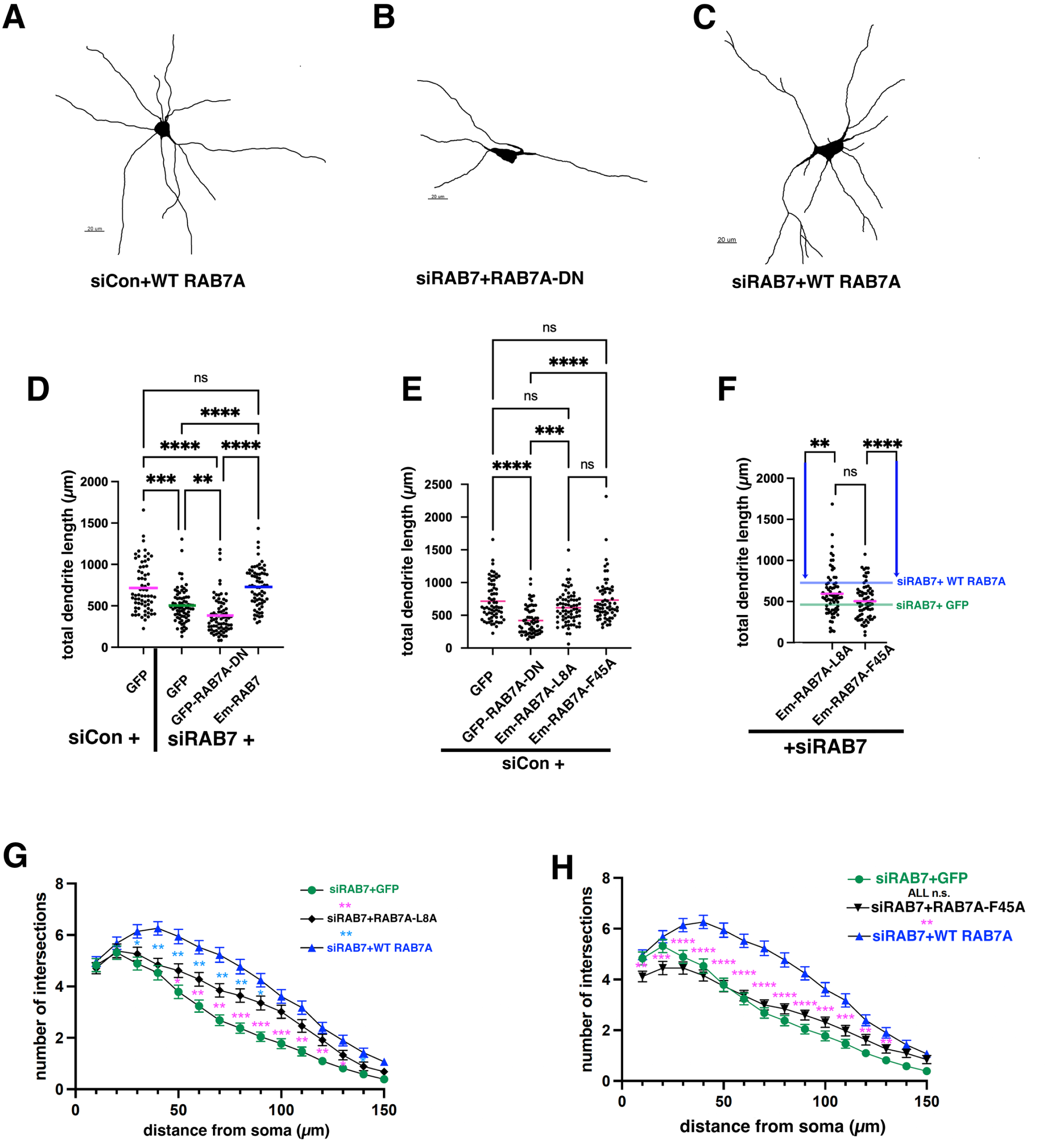
Loss of RILP binding by RAB7A decreases dendrite elaboration. (A-C) Dendrite morphology is simpler in RAB7A depleted neurons and can be rescued by WT RAB7A. Representative tracings are shown for control (A: siCon+WT RAB7A), loss of RAB7A (B: siRAB7+RAB7A-DN), and re-expression of WT RAB7A (C: siRAB7+WT RAB7A). (D-F) DIV5 cultured rat hippocampal neurons were transfected with siControl or siRAB7 and simultaneously with WT or mutant RAB7A for 5 days. Total dendrite length was measured for siCon, siRAB7, siRAB7+RAB7A-DN and siRAB7+WT RAB7A rescue (D). The effect of overexpressing different RAB7A constructs in control neurons (siCon) on total dendrite length was quantified in (E). The ability of RAB7A-L8A and RAB7A-F45A to restore dendrite arborization is quantified in (F). Levels of siRAB7 with and without RAB7A rescue (from data in D) are indicated by blue and green lines, respectively. (G,H) Sholl analysis is shown in (F) for siRAB7 with (blue) or without (green) WT RAB7A co-expression. The rescue ability of RAB7A-L8A is shown in black. Statistics by cell – N = 67-76 cells from 3 independent cultures. Kruskal-Wallis test. ** p<0.01. *** p<0.001. **** p<0.0001.

We additionally analyzed dendrite complexity by Sholl analysis. Because of the large number of conditions, several separate Sholl graphs are shown. Control Sholl analysis (siCon+GFP) is shown in SFig. 3A. siCon+WT RAB7A and siRAB7+WT RAB7A (i.e. rescue) were not statistically different from controls (siCon+GFP). We show a comparison of different RAB7A-depleted conditions in SFig. 3B (siRAB7+GFP vs siCon+RAB7A-DN vs siRAB7+RAB7A-DN). Primary dendrite number (i.e. the number of intersections at the shortest distance measured) was not affected in siRAB7A but reduced when RAB7A-DN is expressed (<4 vs 5 intersections). Consistently, there were fewer branches with increasing distance from the soma for all Rab7A interference conditions (compare SFig. 3A to B). We then compared WT RAB7A rescue to RAB7A-L8A (Fig. 4G) and -F45A (Fig. 4H). Both RAB7A mutants showed significantly reduced rescue ability compared to WT RAB7A, but -F45A showed a stronger phenotype than -L8A and led to a reduction in primary dendrites (Fig. 4H, SFig. 3D). RILP-binding by RAB7A is thus required for normal dendrite elaboration. Interestingly, we see slightly worsened dendrite complexity even in siCon conditions for both RAB7A-L8A and - F45A compared to siCon+GFP (SFig. 3C) especially at shorter distances, even though total dendrite length was not different from control (Fig. 4E). This shows that these mutants have mild dominant-negative effects on dendrites.

### RAB7A-dependent decline in degradative flux is rescued by RAB7A-L8A

We previously showed that inhibition of dynein and downregulation of RAB7A with siRAB7 led to accumulation of the short-lived dendritic membrane receptor NSG2, indicating slowed degradative flux. The accumulation of NSG2 caused by siRAB7 could be rescued by re-expression of WT RAB7A (Yap et al., 2022a). We define degradative flux as the combined multi-step process of endocytosis + maturation + transport + Lys fusion. RILP knockdown in fibroblasts led to deficits in degradative flux (resulting in accumulation of EGFR) (Progida et al., 2007), suggesting the hypothesis that RILP binding by RAB7A is required for normal degradative flux in neurons as well. We thus determined the intensity of DQ-BSA fluorescence, a degradation sensor (Fig. 5). DQ-BSA enters cells by endocytosis and is dye-quenched until it reaches a sufficiently degradative compartment to unquench its red fluorescence (Marwaha and Sharma, 2017). We observed impaired unquenching in siRAB7 (50% of siCon) (Fig. 5A,B,F) which was significantly (but partially) rescued by re-expression of WT RAB7A (back to 80% of siCon; Fig. 5C,F). Rescue by RAB7A-L8A was also partial but significantly less than rescue by WT RAB7A (Fig. 5D,G; 69% of siCon) whereas RAB7A-F45A completely failed to rescue (Fig. 5E,G).

**Figure 5:**
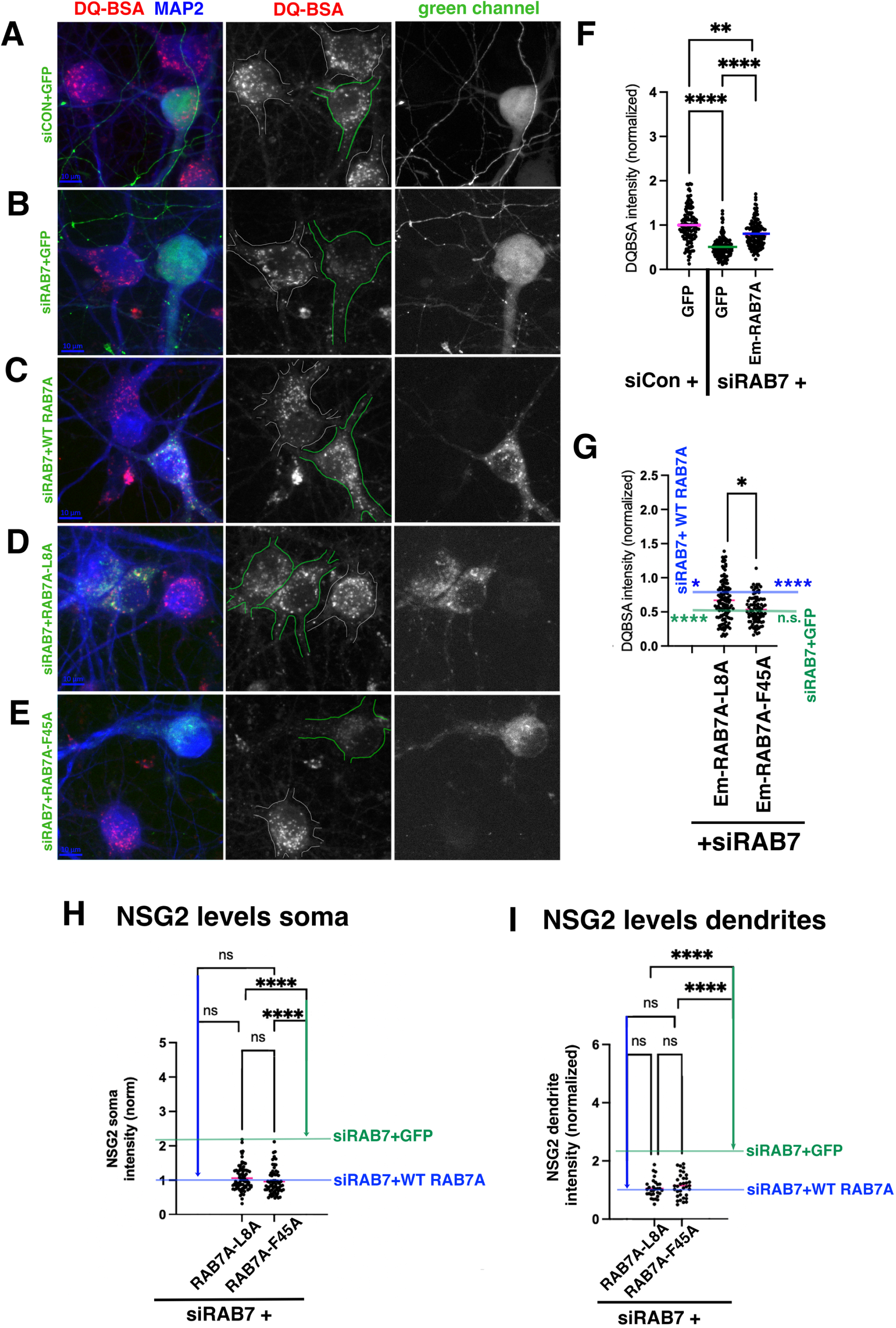
RAB7A-dependent decline in degradative flux is rescued by RAB7A-L8A. (A-G) Degradative capacity (DQ-BSA fluorescence intensity in 4 hours of incubation) was determined for siCon (A), siRAB7 (B) and siRAB7 with re-expression of WT RAB7A (C), RAB7A-L8A (D) and RAB7A-F45A (E). (F,G) Quantification of somatic DQ-BSA intensity in control, siRAB7, and siRAB7+WT RAB7A is shown in (F). The rescue abilities of RAB7A-L8A and RAB7A-F45A compared to siRAB7 (green line) and siRAB7+WT RAB7A rescue (blue line) is shown in (G). Statistics: N = 95-137 cells from 4 independent cultures. Kruskal-Wallis test. ** p<0.01. **** p<0.0001. Note: One extreme high outlier was removed from L8A data set based on Prism outlier testing.sarala(H,I) Levels of endogenous NSG2 in the soma (H) or dendrites (I) of DIV10 hippocampal neurons transfected with siRAB7 and different RAB7A constructs. The levels of WT rescue are indicated by a blue line vs no rescue (siRAB7+GFP; green line). Statistical significance is indicated for each construct with respect to WT rescue (blue arrow) or no rescue (green arrow). Both RAB7A-L8A and RAB7A-F45A completely rescue NSG2 accumulation caused by siRAB7. Statistics: Soma – N = 74-83 cells. Dendrites: N= 30-45 cells (corresponding to 82-118 dendrites). 3 independent cultures. n.s = not significant. **** p<0.0001. Statistics: Kruskal-Wallis.sarala<u>Note</u>: Some of the siRAB7 data (KD, control and WT rescue) were previously published (Yap et al., 2022a) and are re-analyzed here together with the RAB7A mutants which were included in the same set of experiments but not previously published.

Secondly, we analyzed NSG2 levels (Fig. 5H,I) in the soma or dendrites in neurons expressing siRAB7A together with WT RAB7A rescue, RAB7A-L8A or RAB7A-F45A. We previously showed that siRAB7 leads to accumulation of NSG2 in soma and dendrites and can be rescued with WT RAB7A (Yap et al., 2022a). Both RAB7A-L8A or RAB7A-F45A decreased NSG2 accumulation back to WT RAB7A rescue levels (blue line; not statistically significant from siRAB7A+WT RAB7A) in the soma (Fig. 5H) and in dendrites (Fig. 5I). We thus see surprisingly normal NSG2 degradation in RAB7A-L8A expressing neurons.

### RAB7A^fl/fl^ mouse hippocampal neurons in culture

We were surprised that the RILP-binding deficient RAB7A mutants were able to rescue NSG2 degradative flux since they displayed both motility defects (Fig.3) and dendrite morphology defects (Fig.4), and RILP knockdown in fibroblasts impairs EGFR degradation (Progida et al., 2007). Since siRAB7A led to only ∼67% reduction of RAB7A (Fig. 2A), we obtained a RAB7A^fl/fl^ mouse (Roy et al., 2013) in order to more definitively probe the requirement for RILP binding by RAB7A. Hippocampal neurons were cultured from E17 RAB7A^fl/fl^ mouse embryos and transfected with BFP (control) or BFP-Cre plasmid at DIV 5 to excise RAB7A (Fig. 6A). In neurons expressing BFP-Cre, RAB7A levels were over 90% reduced at DIV11 (Fig. 6B). Levels of re-expressed GFP-RAB7A were in a similar range as endogenous RAB7A in most cells (Fig. 6B). About 20% of “rescue” cells (BFP-Cre+GFP-RAB7A) expressed very little RAB7A, and levels were comparable to RAB7A KO (BFP-Cre+GFP). We then repeated the RAB7A-L8A live imaging motility and degradation flux experiments in RAB7A KO neurons in which RAB7A was completely replaced with WT RAB7A (rescue) or with RILP-binding mutants of RAB7A.

**Figure 6:**
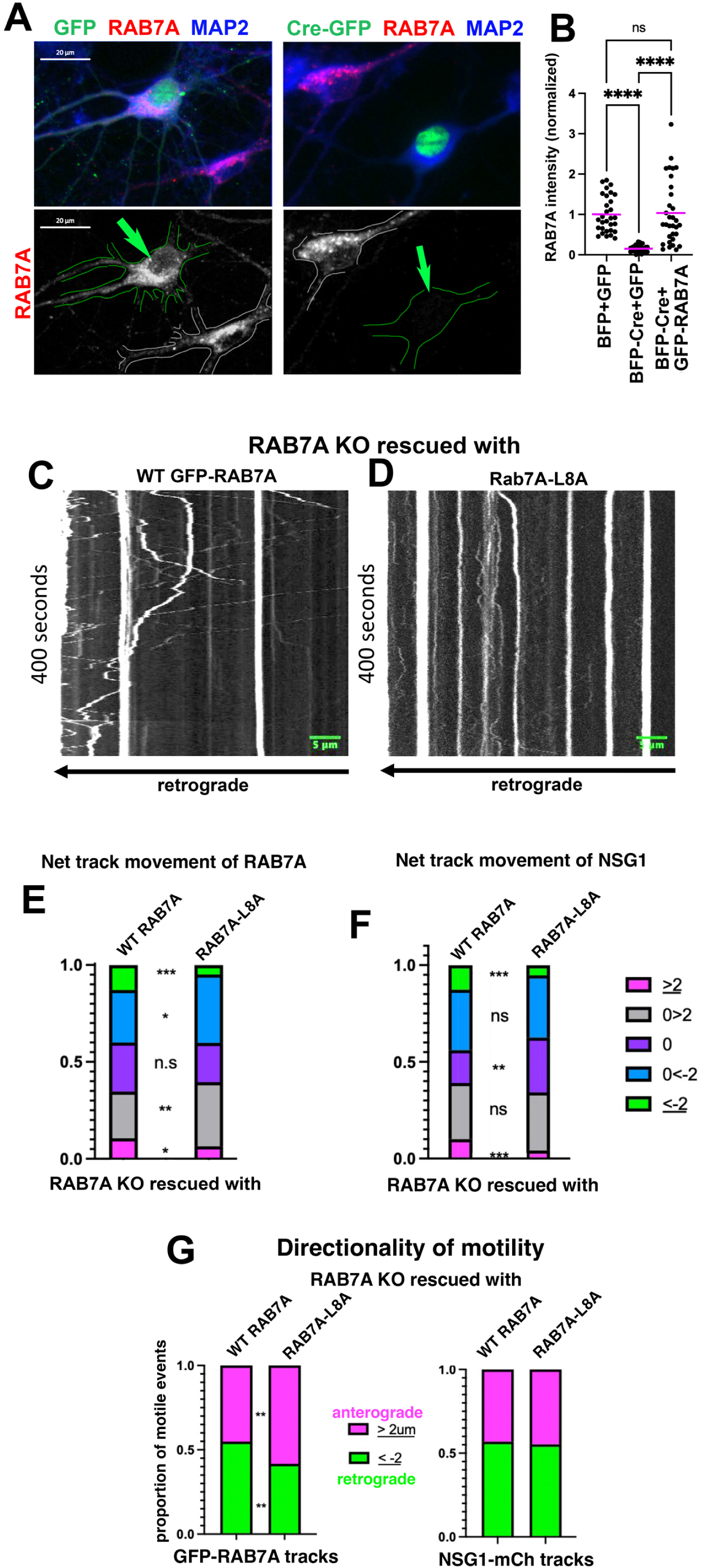
Lack of RILP binding by RAB7A leads to less motility. (A,B) Hippocampal cultures from RAB7A^fl/fl^ mice were transfected at DIV5 with GFP or Cre-GFP for 6 days. (A) Cultures were fixed and stained against MAP2 (A, blue) to identify dendrites and endogenous RAB7A (red). The RAB7A channel is shown alone in the bottom panels in (A). Green arrows point at the cell expressing GFP (left panels) or Cre-GFP (right panels). A non-transfected cell is outlined in white. (B) shows quantification of somatic endogenous RAB7A levels in cells transfected with BFP+GFP (control for transfection), BFP-Cre+GFP (“KO”) or BFP-Cre with WT GFP-RAB7A re-expression (“rescue”). Levels were normalized to BFP+GFP control cells. Median is shown. Statistics – N = 30-32 cells. Kruskal-Wallis test. **** p<0.0001.sarala(C-G) Live imaging of dendritic LEs in RAB7A KO neurons rescued with WT GFP-RAB7A or Em-RAB7A-L8A.sarala(C,D) Representative kymographs of WT GFP-RAB7A (C) and Em-RAB7A-L8A (D) motility are shown in DIV11 RAB7A^fl/fl^ neurons co-transfected with BFP-Cre and NSG1-mCh.saralaRetrograde transport is towards the left. Images were collected for 400sec at a capture rate of 1 frame/sec.sarala(E-G) Kymobutler analysis of kymographs. (E,F) Net track motility of RAB7A (WT or L8A) (E) or of NSG1-Em in WT mCh-RAB7A or mCh-RAB7A-L8A expressing DIV11 hippocampal neurons (F) was binned into 5 categories by directionality (retrograde vs anterograde vs stationary) and extent of movement (small = less than 2 µm in either direction; big = more than 2 µm in either direction), as indicated in the color-coded legend. (G) Only motile (≥ 2 µm) events were included to determine the directionality bias in the motile population for RAB7A tracks (left graph) or NSG1 tracks (right graph). Retrograde bias was decreased by RAB7A-L8A for RAB7A tracks but not for NSG1 tracks. Statistics: N = 24 - 26 cells with 89 and 111 dendrites from two independent cultures. Em-RAB7A tracks: Unpaired t-test. NSG1-mCh tracks: Mann-Whitney. *** p<0.001. **** p<0.0001.

### RAB7A-L8A in RAB7A KO neurons impairs motility of dendritic late endosomes

We carried out live imaging of RAB7A-positive LEs in dendrites of RAB7A-KO neurons re-expressing either WT RAB7A (Fig. 6C) or RAB7A-L8A (Fig. 6D) together with NSG1-mCh. 25.2% (avg) of GFP-RAB7A tracks were stationary (Fig. 6E; purple) and 23.4% (avg) showed >2 µm net displacement in either the anterograde (away from the soma; pink = 10.6% avg) or retrograde (towards the soma; green = 12.8% avg) direction (Fig. 6E). Em-RAB7A-L8A showed significant decrease in both net retrograde (green = 4.8% avg) and anterograde (pink = 6.4% avg) displacement compared to WT RAB7A. When motility of the degradative LE cargo NSG1-mCh was analyzed, it was similarly changed in RAB7A-L8A re-expressing neurons (increased stationary tracks [16.9% to 28.3% avg] and decreased anterograde [10.1% to 4.2% avg] and retrograde tracks [12.7% to 5.3% avg; Fig. 6F). Interestingly, when looking only at the motile population, the proportion of anterograde to retrograde GFP-RAB7A tracks was changed from 55:45 retrograde:anterograde bias in GFP-RAB7A to 42:58 retrograde:anterograde bias in Em-RAB7A-L8A (Fig. 6G). Our results thus show that 53% of the motile RAB7A compartments and 59% of NSG1 compartments rely on RILP binding by RAB7A for directional motility in dendrites. The effect of loss of RILP binding by RAB7A is thus qualitatively the same, but quantitatively larger, in the RAB7A KO compared to siRAB7 (Table 1). We were not able to do live imaging of RAB7A-F45A due to its large cytosolic background which made it impossible to track compartments.

**Table 1:**
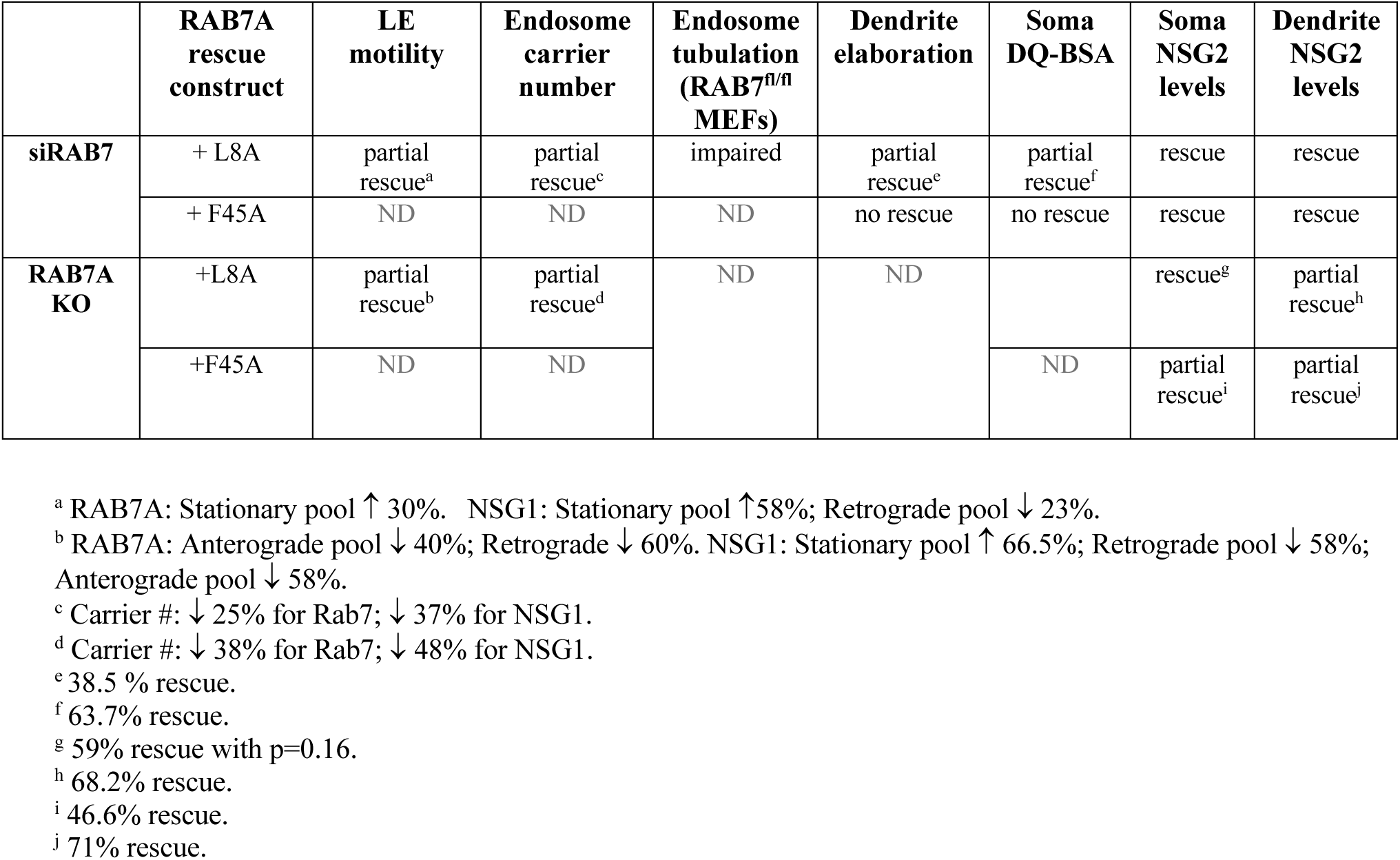
Summary of phenotypes for RAB7A-L8A and RAB7A-F45A.

|  | **RAB7A**<br>**rescue**<br>**construct** | **LE**<br>**motility** | **Endosome**<br>**carrier**<br>**number** | **Endosome**<br>**tubulation**<br>**(RAB7<sup>fl/fl</sup>**<br>**MEFs)** | **Dendrite**<br>**elaboration** | **Soma**<br>**DQ-BSA** | **Soma**<br>**NSG2**<br>**levels** | **Dendrite**<br>**NSG2**<br>**levels** |
| --- | --- | --- | --- | --- | --- | --- | --- | --- |
| **siRAB7** | + L8A | partial<br>rescue<sup>a</sup> | partial<br>rescue<sup>c</sup> | impaired | partial<br>rescue<sup>e</sup> | partial<br>rescue<sup>f</sup> | rescue | rescue |
|  | + F45A | ND | ND | ND | no rescue | no rescue | rescue | rescue |
| **RAB7A**<br>**KO** | +L8A | partial<br>rescue<sup>b</sup> | partial<br>rescue<sup>d</sup> | ND | ND |  | rescue<sup>g</sup> | partial<br>rescue<sup>h</sup> |
|  | +F45A | ND | ND |  |  | ND | partial<br>rescue<sup>i</sup> | partial<br>rescue<sup>j</sup> |

<sup>a</sup> RAB7A: Stationary pool ↑ 30%. NSG1: Stationary pool ↑ 58%; Retrograde pool ↓ 23%.
<sup>b</sup> RAB7A: Anterograde pool ↓ 40%; Retrograde ↓ 60%. NSG1: Stationary pool ↑ 66.5%; Retrograde pool ↓ 58%; Anterograde pool ↓ 58%.
<sup>c</sup> Carrier #: ↓ 25% for Rab7; ↓ 37% for NSG1.
<sup>d</sup> Carrier #: ↓ 38% for Rab7; ↓ 48% for NSG1.
<sup>e</sup> 38.5 % rescue.
<sup>f</sup> 63.7% rescue.
<sup>g</sup> 59% rescue with p=0.16.
<sup>h</sup> 68.2% rescue.
<sup>i</sup> 46.6% rescue.
<sup>j</sup> 71% rescue.

### Lack of RILP binding by RAB7A leads to reduction of carrier numbers and to decreased tubule formation from LEs

During our kymograph analysis, we noticed that the number of scoreable tracks was smaller in movies of RAB7A-L8A expressing neurons, both in the context of siRAB7 (Fig. 7A) and of RAB7A KO (Fig.7 B,C). This was true for both RAB7A tracks (Fig.7 A left side; Fig.7 B) and NSG2 tracks (Fig.7 A right side; Fig.7 C). Since carriers are generated by tubulation and fission from larger LEs (van Weering et al., 2012; Gopaldass et al., 2024), we wondered if we could observe a change in tubulation in RAB7A-L8A. Tubulation is very difficult to observe in neurons, especially in the crowded soma or in narrow dendrites. Therefore, we made MEFs from RAB7A^fl/fl^ embryos (Mulligan et al., 2024), transfected them with RAB7A WT or RAB7A-L8A constructs and then carried out live imaging (Fig.7 D). We find that fewer tubules emanated from Em-RAB7A-L8A compartments compared to WT RAB7A (Fig.7 D red arrows; Fig.7 E). This reduction in LE tubulation likely accounts for the reduced carrier numbers in our live imaging.

**Figure 7:**
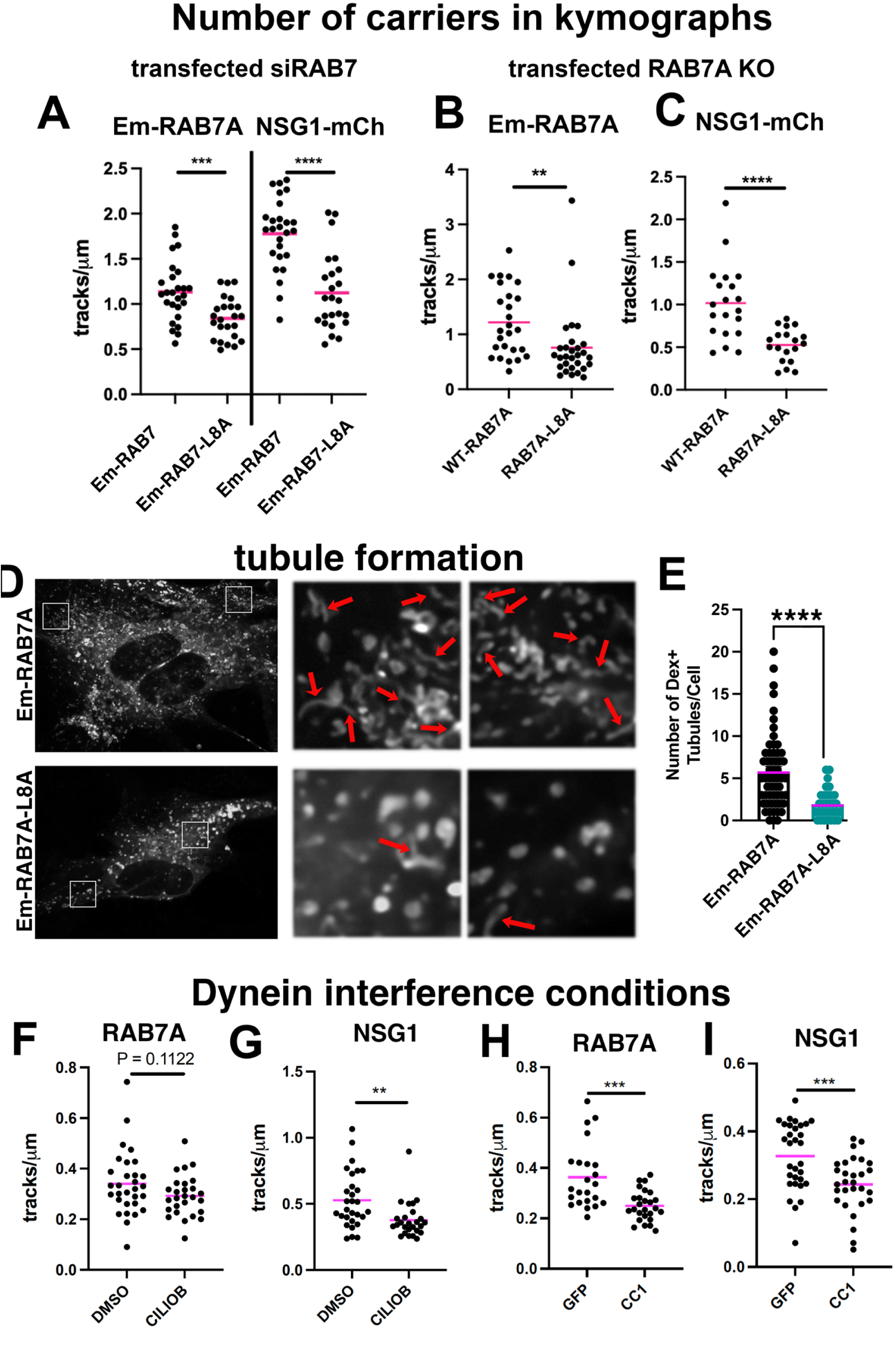
RAB7A-L8A leads to decreased tubule formation from LEs and reduction of carrier numbers. (A) Kymographs analyzed in Fig. 3E-G were used for quantification of track numbers for both GFP/Em-RAB7A positive and NSG1-mCh positive carriers. siRAB7 and NSG1-mCh were co-transfected with either WT RAB7A or RAB7A-L8A and the number of tracks per µm of dendrite was determined using KymoButler. Statistics: N = 24 - 26 cells with 89 and 111 dendrites from two independent cultures. Em-RAB7A tracks: Unpaired t-test. NSG1-mCh tracks: Mann-Whitney. *** p<0.001. **** p<0.0001.sarala(B,C) Number of RAB7A-positive (B) or NSG1-positive (C) carriers/tracks was determined in RAB7A-KO neurons re-expressing WT RAB7A or RAB7A-L8A.saralaStatistics by cell – 1) RAB7A analysis - N = 19-29 neurons (from 2-3 independent cultures representing 84 −124 dendrites). Mann-Whitney or unpaired t-test, as appropriate. * p<0.05, ** p<0.01, *** p<0.001, ****p<0.0001.sarala(D) Live confocal micrographs revealing extent of tubulation in Em-RAB7A WT or EM-RAB7-L8A transfected MEF cells loaded with 100μg/mL tetramethyl-rhodamine dextran (TMRdex) for 2hr. Two regions are shown larger on the right. Tubules = red arrows.sarala(E) Quantification of the number of dually positive Em-Rab7+TMRdex+ tubules/cell. Statistics – Em-RAB7A= 57 cells; EM-RAB7A-L8A= 31 cells. Mann-Whitney-U test. **** p<0.0001.sarala(F-I) Number of carriers in live imaging was reduced by interference with dynein function (re-analysis from data in Yap et al. 2022a). Number of RAB7A-positive (F,H) or NSG1-positive (G,I) carriers/tracks was determined in neurons transfected with mCh-RAB7A or NSG1-mCh separately (for CC1-GFP) or together with Em-RAB7A and NSG1-mCh (for ciliobrevin). Statistics for (F-I) by cell – N= 26-30 cells representing 81-130 dendrites from 3 independent cultures. Mann-Whitney or unpaired t-test, as appropriate. ** p<0.01, *** p<0.001.

Lastly, we reasoned that if tubule formation was dependent of RAB7A-RILP, it should also be dependent on dynein. We thus reanalyzed live imaging data from a previous paper (Yap et al., 2022a) where we inhibited dynein function (Fig. 7F-I). Acute inhibition of dynein motility with ciliobrevin reduced the track number in kymographs of NSG1-positive carriers (Fig.7 G) but did not reach significance for RAB7A carriers (Fig. 7F). Carrier number was reduced for both RAB7A-positive and NSG1-positive carriers in neurons expressing the dynein-interfering construct CC1 (Fig.7 H,I). RILP-binding to RAB7A thus promotes tubulation and carrier formation on LEs in a dynein-dependent manner.

### RAB7A-L8A re-expression in RAB7A KO neurons affects dendritic but not somatic NSG2 degradative flux

We then determined whether WT RAB7A or RAB7A-L8A could rescue degradation flux delays of NSG2 in RAB7A KO neurons. NSG2 was greatly accumulated in soma and dendrites of Cre-GFP expressing RAB7A^fl/fl^ neurons compared to GFP-expressing controls at steady state (Fig. 8A). When RAB7A^fl/fl^ cultures were transfected with GFP or Cre-GFP and then treated with cycloheximide (CHX) for 4 hours to block new translation, NSG2 was largely degraded in GFP controls whereas Cre-GFP expressing RAB7A^fl/fl^ neurons showed high levels of NSG2 remaining in soma and dendrites (Fig. 8B), consistent with blocked degradative flux.

**Figure 8:**
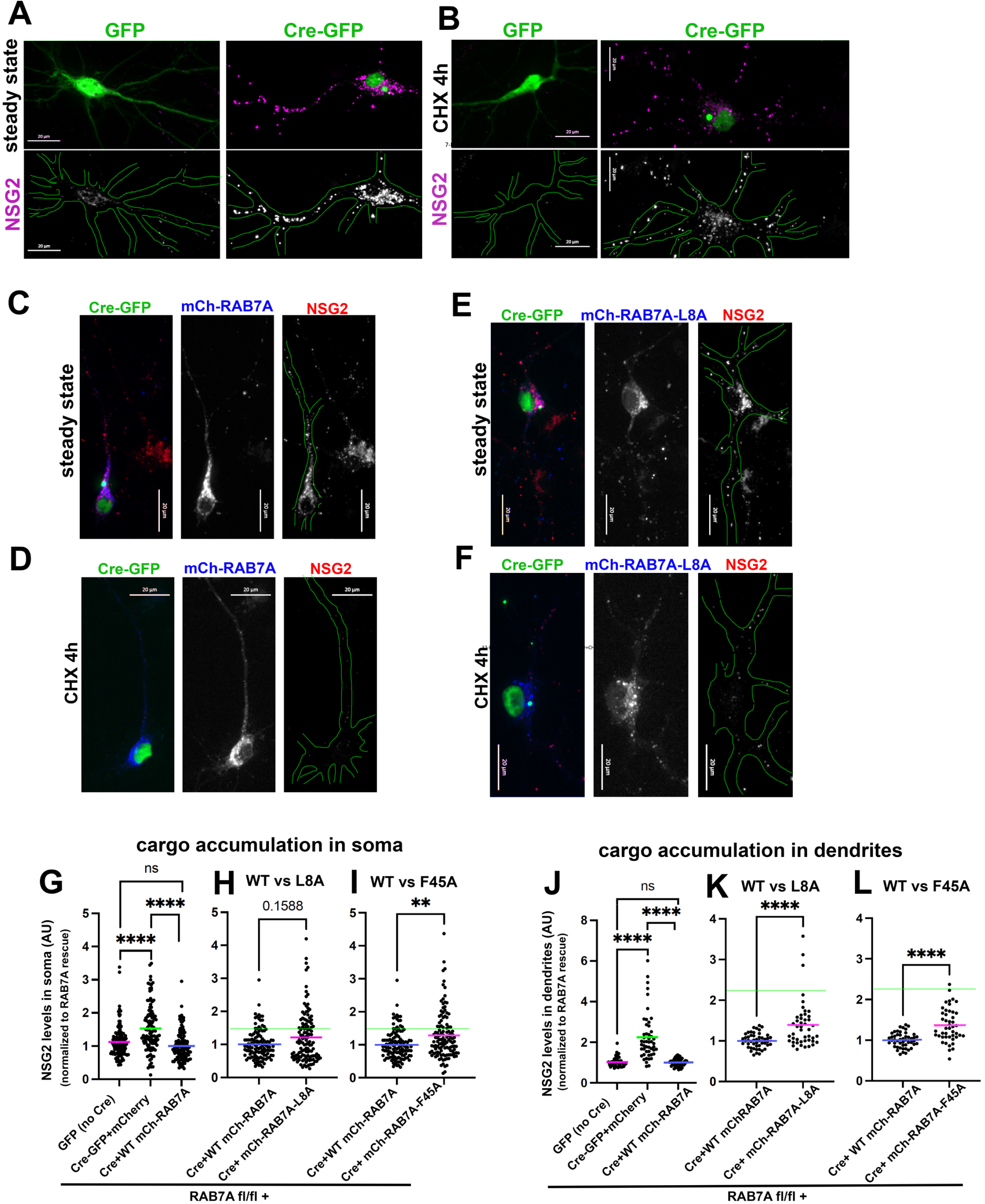
RAB7A-L8A affects dendritic but not somatic NSG2 degradative flux. (A,B) NSG2 degradative flux is impaired in RAB7A KO neurons. DIV5 RAB7A^fl/fl^ hippocampal neurons were transfected with GFP (control; green) or Cre-GFP (RAB7A KO; green) and stained on DIV11 against endogenous NSG2 (magenta) either at steady-state (A) or after 4h of translational block in CHX (B). NSG2 channel alone is shown in the bottom row. MAP2 staining (not shown) was used to create the outline of the transfected cell. NSG2 accumulates in RAB7A KO neurons at steady state (A) and fails to degrade in CHX (B).sarala(C-F) Ability of WT RAB7A and RAB7A-L8A to rescue NSG2 degradation defect are similar. Endogenous NSG2 levels (red) in RAB7A KO neurons transfected with WT mChRAB7A (blue; C,D) or mCh-RAB7A-L8A (blue; E,F) at steady state (C,E) or after 4 h CHX (D,F). Single channels are shown for NSG2 and RAB7A. Cell outlines are based on MAP2 staining. NSG2 cargo accumulation was quantified in the soma (G-I) and dendrites (J-L) for WT RAB7A (G,J), RAB7A-L8A (H,K), and RAB7A-F45A (I,L). Levels were normalized for all graphs to “WT RAB7A rescue”. NSG2 levels for RAB7A KO (i.e. cre-GFP+mCh) is indicated by a green line in (H,I,K,L) for easier comparison.saralaStatistics – soma: (G-I) N = 117-124 cells. --dendrites: (J-L) N = 46-53 cells. 4 independent cultures. G,J: Kruskal-Wallis test. H,I,K,L: unpaired t-test or Mann-Whitney test, as appropriate. ** p<0.01, **** p<0.0001.

Accumulated NSG2 in RAB7A KO neurons could be rescued by re-expression of WT mCh-RAB7A (Fig. 8C; quantified for the soma in Fig. 8G). Similarly, RAB7A KO neurons re-expressing WT RAB7A (“rescue”) showed low NSG2 levels after 4 hours in CHX (Fig. 8D) similar to control neurons (compare to GFP panels in Fig. 8A,B).

We then visualized NSG2 in the soma at steady state (Fig. 8E) or after 4h of CHX chase (Fig. 8F) when RAB7A-L8A was used for the rescue. NSG2 intensity at steady state was rescued by RAB7A-L8A in the soma (Fig. 8E and H) and was not statistically significant from WT RAB7A (Fig. 8H; p=0.16). This observation matches the result from the siRAB7A rescue experiments (Fig. 5H).

When we examined our images, we noticed some bright puncta of NSG2 remaining in RAB7A-L8A. Often these puncta were in dendrites rather than in the soma (Fig. 8F). We therefore quantified NSG2 intensity separately in dendritic endosomes. As we saw before for soma NSG2 intensity, re-expression of WT RAB7A in RAB7A KO neurons rescued dendritic NSG2 accumulation back down to control (GFP) levels (Fig. 8J). Re-expression of RAB7A-L8A, in contrast, did not completely rescue dendritic NSG2 levels (Fig. 8K), indicating that there is a partial requirement for RILP binding by RAB7A for NSG2 removal from dendrites whereas we did not observe such a requirement in the soma (Fig. 8H).

We also determined NSG2 levels when RAB7A-F45A was re-expressed in RAB7A KO neurons compared to WT RAB7A. We see a significant NSG2 accumulation phenotype in the soma (Fig. 8I) as well as in dendrites (Fig. 8L), indicative of partial rescue. A summary of all observed phenotypes is shown in Table 1.

## Discussion

In this work, we dissect the roles of RAB7A-dependent RILP function in neuronal dendrites. RILP is a RAB7A effector implicated in recruitment of dynein to LEs for motility (Cantalupo et al., 2001; Jordens et al., 2001) as well as recruitment of HOPS for fusion with lysosomes (van der Kant et al., 2013; Johansson et al., 2007). In addition, RILP binds to the V1G1 subunit of the vacuolar proton pump (De Luca et al., 2015) and to VPS22, an ESCRT-I component (Wang and Hong, 2006; Progida et al., 2006). RAB7A-RILP binding is thus thought to be required for motility of LEs (via dynein binding), fusion of LEs and autophagosomes with lysosomes (via HOPS binding), as well as potentially the formation of intraluminal vesicles (via ESCRT-I binding) and regulation of acidification (via V1G1). In fact, downregulation of RILP in non-neuronal cell lines leads to accumulation of several LE markers and EGFR degradation defects (Progida et al., 2007). RILP binds HOPS and dynein simultaneously and has been shown to directly link transport and fusion of LEs (van der Kant et al., 2015). RILP might also have RAB7-independent functions both via LC3-binding in axonal autophagosome transport (Khobrekar et al., 2020; but see Cason et al., 2021) and via binding to other RABs (Starling et al., 2016; Kumar et al., 2024; Matsui et al., 2012; Wang and Hong, 2006, 2002). Lastly, axonal retrograde transport of NGF signaling endosomes is reportedly inhibited in sympathetic neurons when RILP is knocked down (Ye et al., 2018; but see caveats in Yap et al., 2023). Because of pervasive off-target problems with RILP knockdown, we use a separation-of-function mutant of RAB7A (RAB7A-L8A) to specifically determine the exact steps where RILP binding by RAB7A is important in dendrites. We find that RILP-binding by RAB7A is required for normal retrograde dendritic motility of LEs, but not for fusion with somatic lysosomes (Fig. 9). Interestingly, normal dendrite morphology requires RILP binding by RAB7A. This indicates that normal dendrite growth/maintenance is dependent on sustained RAB7A/RILP-dependent LE transport (Fig. 9).

**Figure 9:**
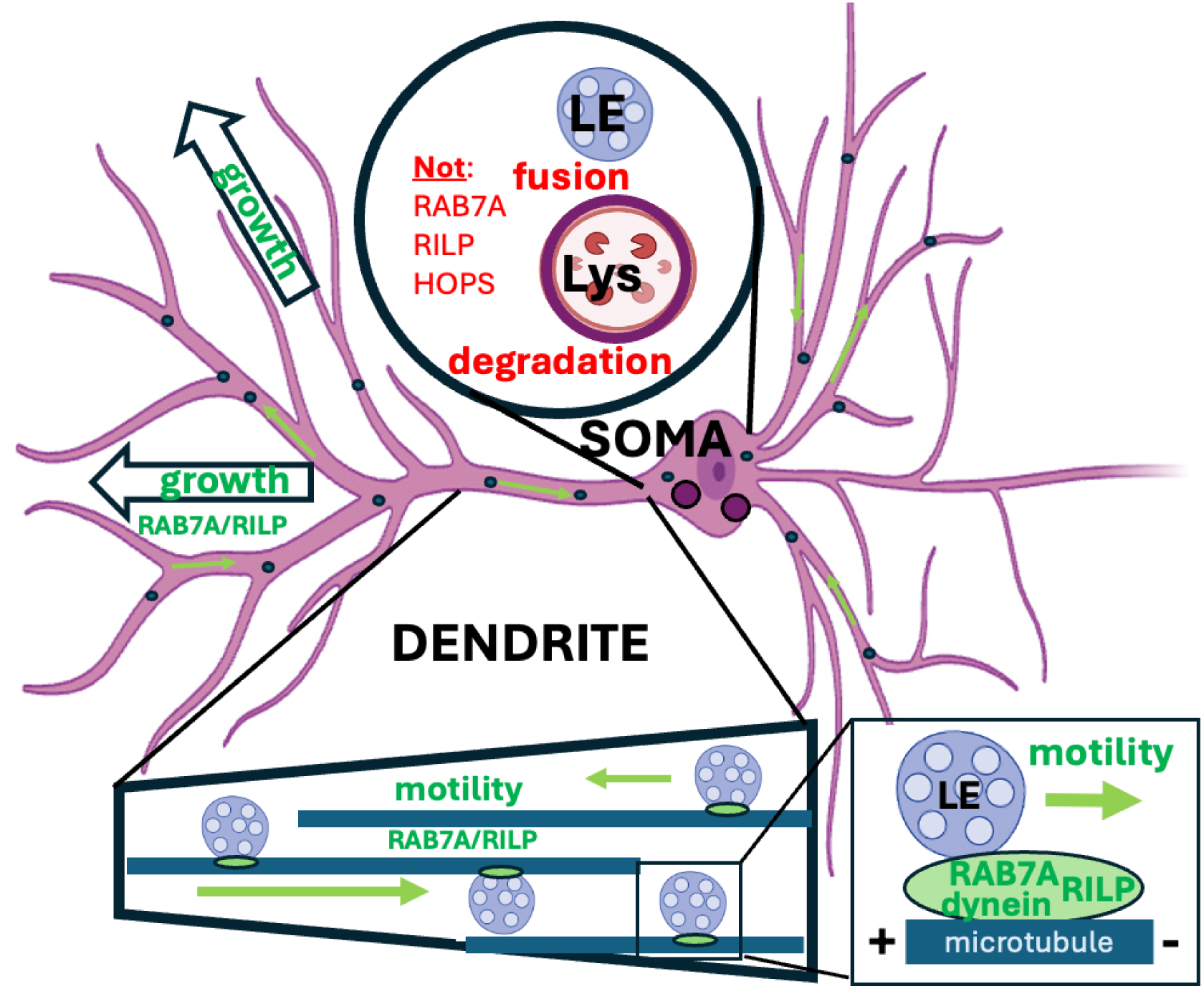
Model for RAB7A-RILP dependent and independent steps in dendrites: RAB7A-RILP is required for normal LE motility and dendrite growth but not for fusion with somatic lysosomes and degradation. RAB7A-RILP-dynein promote LE motility (green arrows) in dendrites. Fusion with somatic lysosomes which leads to subsequent degradation (red) does not require RAB7A-RILP but might be dependent on other proteins. Elaboration of a normal dendritic arbor depends on RAB7A-RILP function (green “growth” arrows), implicating transport of LEs in dendrites in promoting dendrite growth.

### Roles for RILP-RAB7A in LE motility and carrier formation

Our results show that RILP binding by RAB7A contributes significantly to retrograde motility of LEs in dendrites (see Table 1), but there is residual motility in RAB7A-L8A expressing neurons of a subset of LEs. This residual motility could be due to RAB7A-FYCO1-kinesin complexes (Pankiv et al., 2010) moving LEs towards the soma on minus-end out microtubules. Alternatively, this subset of LEs could link to motors independently of RAB7A, (Marwaha et al., 2017). Loss of RILP binding by RAB7A also reduces the number of motile carriers in dendrites. In non-neuronal cells, we observe reduction of LE tubulation which is a necessary step for carrier formation. This finding is consistent with carrier formation requiring microtubule motors (Gopaldass et al., 2024), in our case dynein. Due to the dual reduction in carrier formation and subsequent retrograde transport to somatic lysosomes, dendritic cargos accumulate in LEs in RAB7A-L8A expressing neurons. We conclude that RILP is in fact the endogenous dynein adaptor for a substantial and functionally important set of LEs to support carrier formation and retrograde transport.

### RILP binding by RAB7A is not required for somatic degradation of dendritic cargos

HOPS is a tethering complex required for LE-LE and LE-Lys fusion (Pols et al., 2013; Wartosch et al., 2015; van der Beek et al., 2024). RAB7A binds to and stabilizes HOPS/RILP on LEs (Lin et al., 2014), and knockdown of RILP in fibroblasts leads to failure to degrade EGFR (Progida et al., 2007). We thus expected to see failure of LEs to fuse with lysosomes in RAB7A-L8A expressing neurons resulting in degradation block. Surprisingly, we did not observe this. Degradative activity in the soma is not impaired by RAB7A-L8A, suggesting that fusion of LEs with degradative lysosomes does not require RILP binding by RAB7A. Several alternative pathways for LE-Lys fusion have been described (Jongsma et al., 2020; Schleinitz et al., 2023; Marwaha et al., 2017), all of which would still be operative and could sustain fusion in the absence of RILP-RAB7A binding. In fact, in overexpression experiments, RAB7A-L8A still associates with one of the alternative adaptors PLEKHM1 (CCY, unpublished observations).

Initial reports suggested that RAB7A in a complex with RILP-HOPS-dynein-ORP1L coordinates transport with fusion (van der Kant et al., 2013) but more recent reports suggest that RAB2A rather than RAB7A provides the HOPS tethering activity (Schleinitz et al., 2023). Our findings are consistent with the latter model. Since RAB7A-L8A maintains weak, RILP-independent HOPS binding (Lin et al., 2014) as well as possibly being still able to bind HOPS via a different adaptor (i.e. SKIP) (Jongsma et al., 2020), it can likely still aid in the HOPS handoff even in the absence of RILP binding. In addition, RILP can also bind other RAB proteins, RAB34 and RAB36 (Wang and Hong, 2006; Colucci et al., 2005; Kumar et al., 2024; Starling et al., 2016), with less well understood roles in endo-lysosomal function. These RABs could provide redundant RILP-dependent functions in the soma since our data clearly show that somatic degradation does not require RAB7A-RILP binding.

The lack of a somatic degradation phenotype for NSG2 and the partial rescue of DQ-BSA fluorescence also strongly suggest that lysosomes are fundamentally functional in the absence of RAB7A-RILP binding. RAB7A-RILP-V1G1 binding (De Luca et al., 2015) is thus not required for lysosome acidification. In fact, we recently discovered that V1G1 acts upstream of RAB7A-RILP rather than downstream (Mulligan et al., 2024).

### RAB7A-F45A has more severe phenotypes than RAB7A-L8A

RAB7A-L8A and RAB7A-F45A share several phenotypes but RAB7A-F45A phenotypes are often more severe than RAB7A-L8A (Table 1). This is not surprising since F45 is part of the hydrophobic triad (Wu et al., 2005; Khan et al., 2021) in the interswitch region of the RAB7A conformational sensor whereas L8 lies at the extreme N-terminus. Interestingly, modeling of RAB7A binding to its effector PDZD8 shows a direct contact with F45 (Khan et al., 2021). It is thus likely that RAB7A-F45A has lower binding to multiple other RAB7A effectors in addition to RILP. This manifests in our experiments by increased cytosolic localization of RAB7A-F45A with reduced recruitment to FYCO1-stabilized compartments. It is actually surprising that RAB7A-F45A can still rescue some siRAB7A phenotypes and partially rescue some RAB7A KO phenotypes (Table 1). We find that binding to some effectors is still mostly intact (such as to VPS35) likely accounting for the increased rescue ability of RAB7A-F45A compared to the dominant-negative RAB7A-T22N which binds none of its effectors. RAB7A-L8A is thus the more stringent separation-of-function mutant for RILP binding.

### RAB7A-RILP binding supports dendrite growth/maintenance

RILP-binding by RAB7A is also required for normal dendrite morphology. We find it notable that the observed dendrite morphology phenotypes in siRAB7/RAB7A-L8A occur independently of degradation blocks since dysfunctional proteostasis is often viewed as the culprit for poor neuronal morphology and health (Lottes and Cox, 2020). We propose that the observed dendrite phenotypes reflect a requirement for normal motility of LEs in dendrites unrelated to degradative flux per se. We speculate that dendrite defects arise due to trafficking defects of one or more cargos required for dendrite growth. For example, it is known that multiple growth factors can stimulate dendrite growth (Valnegri et al., 2015), often dependent on endocytosis of the ligand-bound receptors which are transported in and signal from endosomes, (Cosker and Segal, 2014; Yamashita and Kuruvilla, 2016; Lehigh et al., 2017). Trafficking of neurotrophin signaling endosomes in dendrites might thus require RAB7A-RILP. The need for RAB7A late endosomes is well established in axonal neurotrophic signaling by NGF-TrkA (Ye et al., 2018) and has been linked to dendritic growth (Moya-Alvarado et al., 2023). Roles for RAB7A in signaling endosome function in dendrites is not well understood. In fact, BDNF-TrkB signaling for dendrite growth has been mostly linked to RAB11-positive recycling endosomes rather than RAB7A-positive LEs (González-Gutiérrez et al., 2020; Moya-Alvarado et al., 2018; Lazo et al., 2013). Understanding potential defects on dendritic signaling endosome dynamics in the context of RAB7A-L8A will be an interesting question to tackle in future work.

## Conclusion

Our work establishes endogenous RILP as a functional RAB7A-dependent LE adaptor for dynein in dendrites and demonstrates a requirement for RAB7A-RILP dependent dendritic LE motility in dendrite arborization independent of degradation (Fig.9).

## Materials and Methods

*<u>Reagents</u>* used are listed in the Table below.

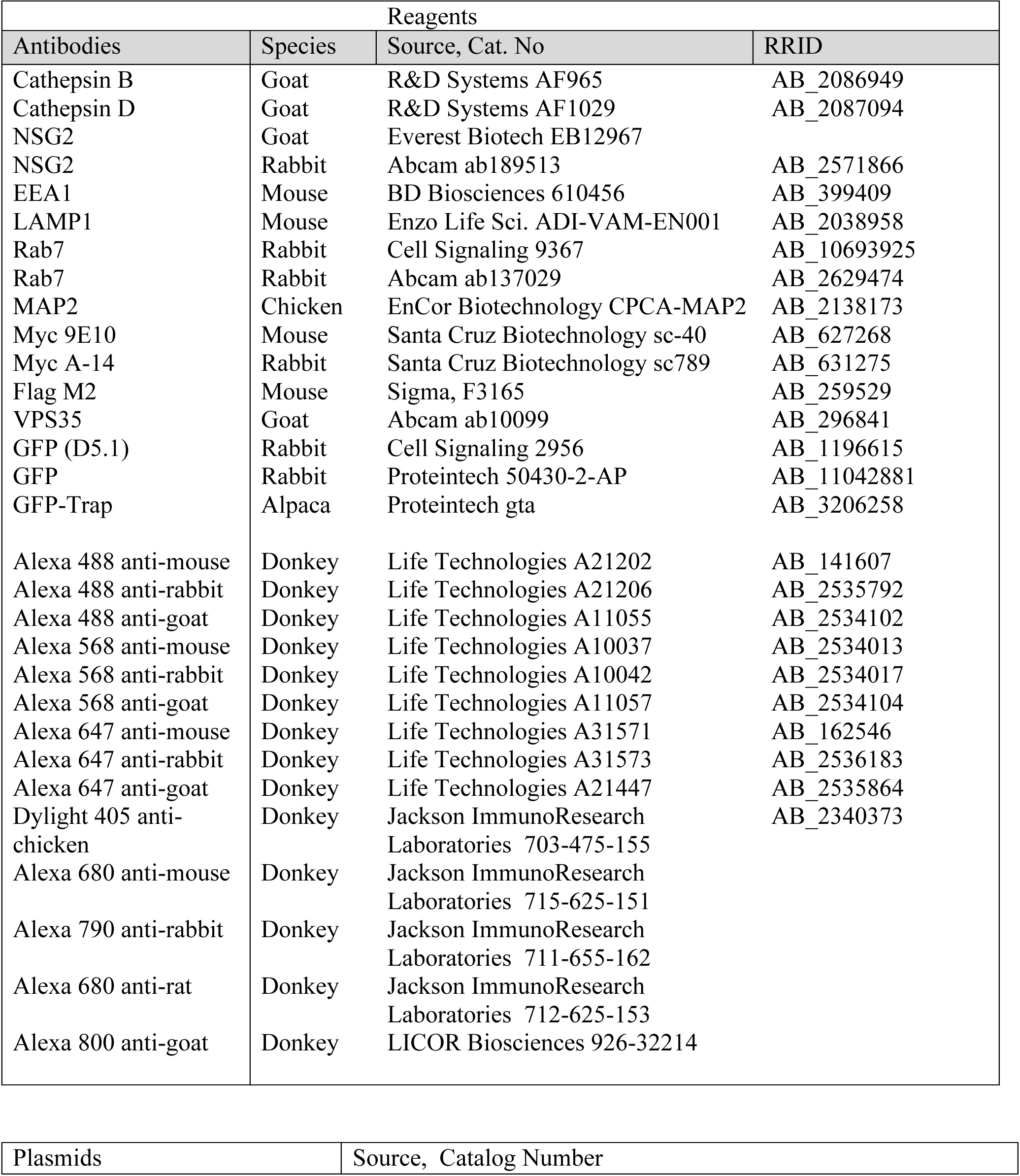
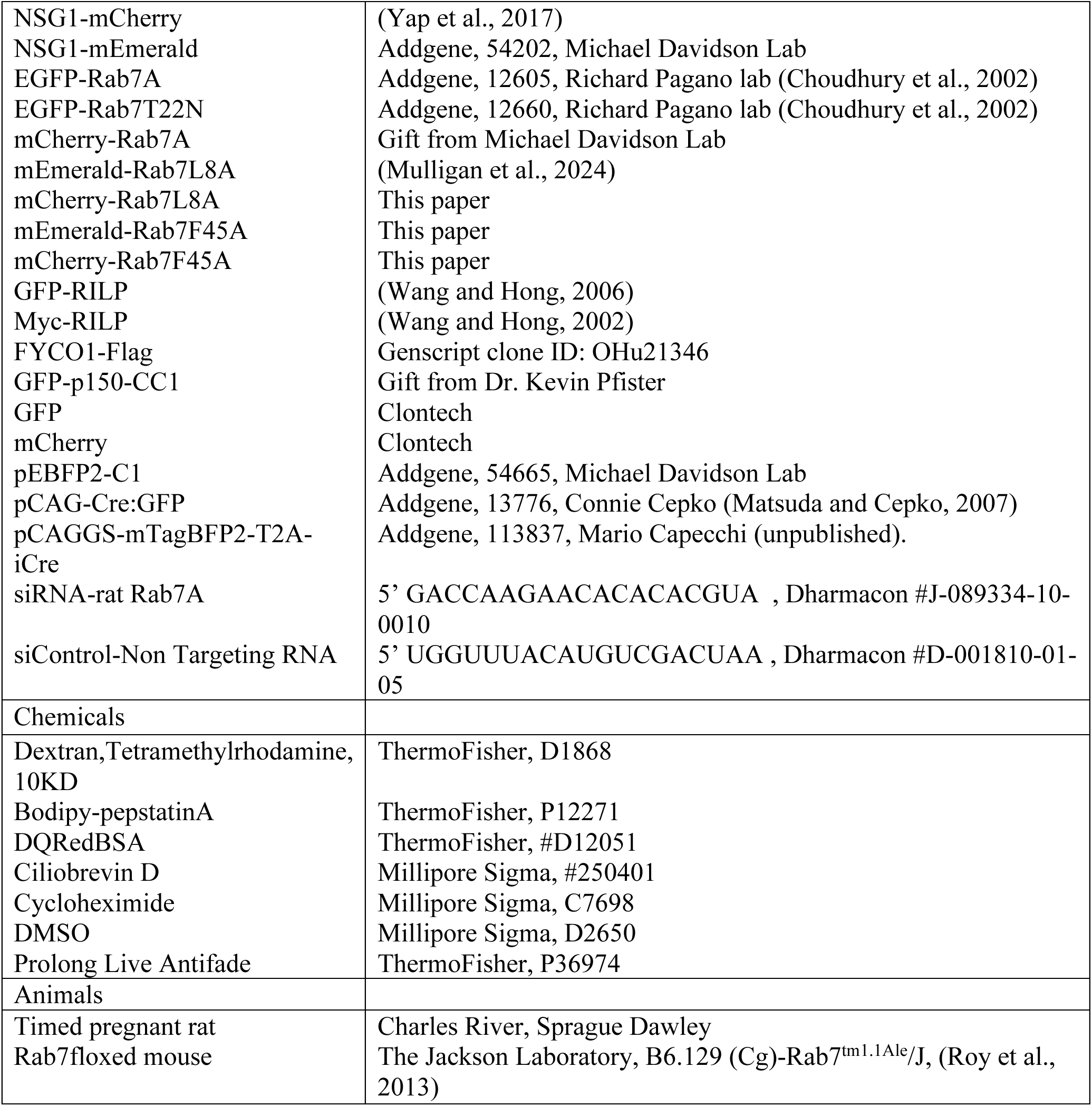

### Plasmid Construction

As described previously (Mulligan et al., 2024), mEmerald-Rab7A-L8A and mCherry-Rab7A-L8A were generated using gene synthesis with a point mutation at residue 8 from leucine to alanine (L8A) in human Rab7A gene and cloned into pEmerald-C1 (Addgene #54734) or pmCherry-C1 at XhoI-BamHI sites by Genscript. Similarly, mEmerald-Rab7A-F45A and mCherry-Rab7A-F45A were generated with a point mutation at residue 45 from phenylalanine to alanine (F45A) and cloned into pmEmerald-C1 and pmCherry-C1 at XhoI-BamHI sites, respectively.

### Neuronal cultures and transfection

Neuronal cultures were prepared as described (Yap et al., 2022a). In brief, the cultures were prepared from either E18 rat hippocampi, or E16.5 Rab7^fl/fl^ mouse hippocampi as approved by the University of Virginia Animal Care and Use Committee. All experiments were performed in accordance with relevant guidelines and regulations (ACUC protocol #3422). Hippocampi from all embryos in one litter were combined and thus contained male and female animals. Cells were plated on poly-L-lysine coated coverslips and incubated with plating medium containing DMEM medium with 10% horse serum. For live imaging use, neurons were plated on a 35mm glass bottom microwell dish (MatTek). After 4 h, the plating medium was removed and replaced with serum-free medium supplemented with B27 (ThermoFisher), and neurons were cultured for 5–10 DIV (days *in vitro*) for experimental use.

Transfections were carried out using Lipofectamine 2000 (Invitrogen). Neurons at DIV7-8 were transfected with either GFP-RILP, myc-RILP, GFP-RAB7A plus myc-RILP, GFP-RAB7A plus FYCO1-FLAG, Em-RAB7A-L8A plus FYCO1-Flag, Em-RAB7A-F45A plus FYCO1-Flag, or GFP-RAB7A-T22N plus FYCO1-FLAG for 40 hours or short (8 hours) for experiments that studied the localization of RILP. For RAB7A knockdown experiments, neurons were transfected with previously validated siRNA against rat RAB7A or siControl/non-targeting RNA (as in (Yap et al., 2022a)). Depending on the experiment, GFP plasmid vector, GFP-RAB7A, GFP-RAB7A-DN, mEmerald-RAB7A-L8A or mEmerald-RAB7A-F45A were co-transfected along with siRAB7 at DIV5 and incubated for 5-6 more days. To determine the RAB7A protein level in RAB7A knockout neurons, mouse RAB7A^fl/fl^ hippocampal neurons were transfected at DIV5 with either GFP, cre-GFP, BFP plus GFP, BFP-cre plus GFP, or BFP-cre plus GFP-RAB7A for 6 days. To investigate the effects of RAB7A knockout on NSG2 degradation, hippocampal neurons at DIV5 were transfected with either GFP, cre-GFP, cre-GFP plus mCherry-RAB7A, cre-GFP plus mCherry-RAB7A-L8A, or cre-GFP plus mCherry-RAB7A-F45A for 6 days, followed by incubation with/without cycloheximide (CHX, 20µg/ml) for 4hrs. All transfection experiments were repeated in at least 2-3 independently derived cultures.

### Cell lines

293 cells were maintained in DMEM plus 10% fetal bovine serum without antibiotic to use for immunoprecipitations. Primary mouse fibroblasts (MEF) were isolated from Rab7^fl/fl^ mouse embryos and maintained as described in (Mulligan et al., 2024).

### Live Imaging

Live imaging was performed as described previously in (Yap et al., 2018, 2017). In brief, rat or mouse RAB7A^fl/fl^ neurons were transfected at DIV5/6 with the following plasmid combinations: siRAB7 with either NSG1-mCherry plus GFP-RAB7A or with NSG1-mCherry plus mEmerald RAB7A-L8A, cre-BFP with either NSG1-mCherry plus GFP-RAB7A, NSG1-mCherry plus mEmerald-RAB7A-L8A, NSG1-mEmerald plus mCherry-RAB7A, or NSG1-mEmerald plus mCherry-RAB7A-L8A for 6 days. To investigate the effects of cytoplasmic dynein activity on endosome carrier formation, live imaging reported previously in (Yap et al., 2022) was re-analyzed. Briefly, rat neurons were transfected at DIV7/8 for 40 hours with the following plasmids: GFP plus mCherry-Rab7A, GFP-p150-CC1 plus mCherry-Rab7A, GFP plus NSG1-mCherry, and GFP-p150-CC1 plus NSG1-mCherry. For acute inhibition of cytoplasmic dynein activity, rat neurons were transfected with either GFP-Rab7 or NSG1-mCherry and treated with 50 µM of ciliobrevin D or DMSO 90 minutes prior to imaging (Yap et al., 2022). Neurons were maintained in Phenol Red-free Neurobasal medium and Prolong Live anti-fade (Life Technologies) was added 30 minutes before live imaging. All live imaging was performed either on a 37°C controlled Nikon TE 2000 microscope using a 60×/NA1.45 oil objective, equipped with a Yokogawa CSU 10 spinning disc, or, on a 37°C heated stage in a chamber with 5% CO_2_ on an inverted Zeiss LSM880 confocal microscope using a 40× water objective (LD-C Apochromat 1.2W). All dual/trio channels images were acquired simultaneously with a 512×512 Hamamatsu 9100c-13 EM-BT camera. Images were captured at a rate of one image every second for 400 frames with perfect focus on. Laser lines at 488 nm for GFP/Emerald, at 405nm for BFP and at 594 nm for mCherry expression were used. Live imaging for any given set of transfected constructs was repeated in at least 2-3 independent cultures. Each time, 6-8 cells per transfected construct were imaged live. All dendrites on all cells were included in the analysis and none were excluded.

### Immunocytochemistry

Immunostaining of neurons was carried out as described (Yap et al., 2017). Neurons were fixed in 2% paraformaldehyde/4% sucrose/PBS in 50% conditioned medium at room temperature for 30 minutes, quenched in 10 mM glycine/PBS for 10 minutes. After washing with PBS, cells were then blocked in 5% horse serum/1% BSA/PBS ± 0.2% TritonX-100 or 0.1% saponin for 20 minutes. All antibodies were diluted in 1% BSA/PBS and incubated for 1 hour. Coverslips were mounted in Prolong Gold mounting medium (ThermoFisher, P36934) and viewed on a Zeiss Z1-Observer with a 40x objective (EC Plan-Neofluar 40x/0.9 Pol WD = 0.41). Apotome structured illumination was used for most images. Images were captured with the Axiocam503 camera using Zen software (Zeiss) and processed identically in Adobe Photoshop. No non-linear image adjustments were performed.

### Bodipy-pepstatinA labeling

Bodipy FL covalently conjugated Pepstatin A is an inhibitor of cathepsin D and binds active cathepsin at pH4.5 in live cells. Thus, it is used to live label acidified, degradative compartments using the manufacturer’s protocol. In brief, transfected neurons were incubated live with 1μM Bodipy-PepstatinA for 45 min, followed by 3 washes with PBS and then fixed with PFA at 2% final concentration at room temperature for 30 minutes. Immunostaining was conducted as described above.

### DQBSA labeling for measurement of degradative flux

To measure the degradative flux, transfected neurons were incubated with 50ug/ml of DQBSA for 4 hours, followed by 3 washes with PBS and then fixed with 2% PFA at room temperature for 30 minutes. Immunostaining was performed as described above.

### Dextran Tubulation

As described in (Mulligan et al., 2024), MEFs transfected with either mEmerald-RAB7A or mEmerald-RAB7-L8A were live labeled with 10K TMR-dextran at 100 μg/ml for 2hrs. After 1X washing with PBS, the cells were live imaged in phenol-red free complete DMEM medium on a 37°C heated chamber on Nikon confocal microscope (Ti2-E with AX-R) using a 60X/NA1.45 oil objective.

### Co-immunoprecipitation and Western Blot

293 cells were transfected with the following plasmids combination: GFP-RAB7A plus myc-RILP, Em-RAB7-L8A plus myc-RILP, Em-RAB7A-F45A plus myc-RILP, GFP-RAB7A-T22N plus myc-RILP. All transfections were conducted using Lipofectamine 2000 (Invitrogen) according to the manufacturer’s protocol. 48 hrs after transfection, HEK293 cells were lysed for 45 min on ice in buffer containing 20 mm Tris-HCl (pH 7.4), 2 mm EDTA, 2 mm EGTA, 0.1 mm DTT, 100 mm KCl, 10 mm NaF, 1 mm PMSF, 1% Triton X-100 and Complete Protease Inhibitor Cocktail. Lysed cells were centrifuged at 12,000 rpm for 15 min, and the supernatants were then precleared with protein G Sepharose (GE Healthcare) at 4°C for 3 h. Precleared lysates were immunoprecipitated with either GFP-Trap or RFP-Trap overnight at 4°C. Immunoprecipitates were washed 5× with lysis buffer containing 0.1% Triton X-100 and eluted by 150 mm Tris-HCl (pH 6.8), 2% SDS, 5% β-mercapto-ethanol, and 8% glycerol at 95°C. Samples were subjected to SDS-PAGE and Western blot analysis as described previously (Mulligan et al., 2024; Bott et al., 2020; Yap et al., 2018). In brief, protein samples were run on a 4%–20% gradient polyacrylamide gel. Proteins were transferred onto nitrocellulose with the Bio-Rad Transblot Turbo according to the manufacturer’s protocol. Membrane blocking was performed in Li-Cor TBS blocking buffer for 1 h at room temperature. Primary and secondary antibodies were diluted in 50% blocking buffer, 50% TBS, and 0.1% Tween-20. Membranes were incubated with primary antibodies for overnight at 4°C, followed by near-infrared secondary antibodies for 1 h at room temperature. All washes were done in TBS+0.1% Tween 20. Blots were imaged with the dual-color Li-Cor Odyssey CLx imager with LI-COR ImageStudio software.

### Quantification of RAB7A soma intensity (Knockdown and Knockout)

Soma intensities were quantified using Imaris 9.5.1. Briefly, the transfected cells were identified, the somata were masked and the average intensity was measured after background correction. Between 29-46 cells per experiment were quantified.

### Kymograph Analysis

All kymographs were generated using the Multi-Kymograph plug-in for FIJI as described (Digilio et al., 2023). Kymographs were analyzed using the machine learning program Kymobutler (Jakobs et al., 2019). Net anterograde and retro-grade track distances are reported regardless of pauses or direction changes during the trajectory, as in (Yap et al., 2022a).

### Measurement of dendrite length

All dendrites on each transfected cell were traced, and their lengths measured using Filament Tracer in Imaris 9.5.1, as in (Yap et al., 2022a). Total length per cell is reported for 62-73 cells/condition from three independent experiments. Sholl calculations were done within Imaris using the same filament tracings.

### Quantification of RAB7A compartment intensity on FYCO1 compartments

Somata of transfected cells were masked using Imaris 9.5.1. Within the masked regions, surfaces representing FYCO1 compartments were created and the sum intensity of RAB7A fluorescence reported for co-localization with the FYCO1 mask vs total RAB7A fluorescence in the soma. The ratio of FYCO1-associated vs cytosolic intensities was determined. N = 30-38 cells from two separate experiments were analyzed.

### Quantification of intensity of NSG2 in dendrites

Because of the extensive crisscrossing of dendrites from different neurons in the field of view, it is difficult to determine the total NSG2 fluorescence in dendrites, and we use an object-based methodology for dendrites. Transfected cells were identified and non-crossing dendrites were traced straightened using FIJI. Spot objects were created for labeled markers within the straightened dendrites. We determined the average NSG2 intensity/endosome using the “Spot” function of Imaris 9.5.1. to identify individual endosomes. Data were normalized to the RAB7A rescue, reported by cell and combined from two to three independent experiments. Twelve to twenty-five cells with 1-5 dendrites/cell per experiment were counted.

### Reanalysis of Data

We reanalyzed two sets of data (Yap et al. 2022a). The first set was NSG2 levels after siRAB7 adding the RAB7A mutants to the previous analysis. The RAB7A mutants were run at the same time as the other conditions but were not published in the previous paper. The second set is the live imaging data of dynein interference where track numbers were not analyzed.

### Experimental Design and Statistical Analyses

All experiments were performed with all conditions in parallel in the same cultures at the same time. This includes live imaging of all plasmid combinations for any given experiment on the same day. Therefore, different plasmids transfections can be compared with more confidence since culture-to-culture variation would affect all experimental conditions equally on any given day. All experiments were repeated several times in independent cultures, as indicated in the figure legends. All data were analyzed using Prism software version 10.3.1. Each dataset was first evaluated for normality by using the Shapiro-Wilk normality test. The result was used to determine whether parametric or nonparametric tests were used. When more than one comparison was made, the corresponding ANOVA test was used. When only two conditions were compared, a t-test was used. Statistical tests are indicated in the figure legends. All N values are shown below.

### Number of cells quantified for each graph

Fig. 2A

siCon+GFP=33 cells; siRAB7+GFP=29 cells; siRAB7+WT RAB7A=46 cells; siRAB7+RAB7A-L8A=33 cells; siRAB7+RAB7A-F45A=31 cells.

Fig. 2J

WT RAB7A=38 cells, RAB7A-L8A=35 cells, RAB7A-F45A=30 cells, DN-RAB7A=38 cells from 2 independent experiments.

Fig. 3C

WT RAB7A: N = 26 neurons (from 2 independent cultures representing 111 dendrites). RAB7A-L8A: N = 24 neurons (from 2 independent cultures representing 89 dendrites).

Fig. 4G,H

siCon+GFP=67 cells, siRAB7+GFP=73 cells, siRAB7+RAB7A-DN=76 cells, siRAB7A+WT RAB7A=69 cells, siRAB7A+RAB7A-L8A=72 cells, siRAB7A+RAB7A-F45A=68 cells from 3 independent cultures.

Fig. 5F,G

siCon+GFP=132 cells, siRAB7+GFP=133 cells, siRAB7A+WT RAB7A=137 cells, siRAB7A+RAB7A-L8A=127 cells, siRAB7A+RAB7A-F45A=95 cells from 4 independent cultures.

Fig. 5H,I

Soma – siRAB7+WT RAB7A = 74 cells; siRAB7+RAB7A-DN=85 cells; siRAB7+RAB7A-L8A=77 cells; siRAB7+RAB7A-F45A=69 cells; siRAB7+GFP=83 cells.

Dendrites: siRAB7+GFP=45 cells (118 dendrites); siRAB7+WT RAB7A=30 cells (96 dendrites); siRAB7+RAB7A-L8A=30 cells (82 dendrites); siRAB7+RAB7A-F45A=34 cells (91 dendrites); siRAB7+RAB7-DN=40 cells (92 dendrites). 3 independent cultures.

Fig. 6B

BFP+GFP=30 cells; Cre-BFP+GFP=32 cells; Cre-BFP+WT GFP-RAB7A=32 cells.

Fig. 6G and Fig. 7A

WT RAB7A = 26 cells with 111 dendrites; RAB7A-L8A = 24 cells with 89 dendrites from two independent cultures.

Fig. 7B,C

RAB7A analysis –

WT RAB7A: N = 26 neurons (from 3 independent cultures representing 124 dendrites).

RAB7A-L8A: N = 29 neurons (from 3 independent cultures representing 112 dendrites).

NSG1 analysis –

WT RAB7A: N = 20 neurons (from 2 independent cultures representing 84 dendrites).

RAB7A-L8A: N = 19 neurons (from 2 independent cultures representing 71 dendrites).

Fig. 7F-I

ciliobrevin experiments –

DMSO control: N = 29 cells (from 3 independent cultures representing 128 dendrites).

CILIOB: N = 26 cells (from 3 independent cultures representing 121 dendrites).

CC1-GFP experiments –

GFP control: NSG1 31 cells (from 3 independent cultures representing 121 dendrites) and RAB7A 23 cells (from 3 independent cultures representing 81 dendrites).

CC1-GFP: NSG1 29 cells (from 3 independent cultures representing 110 dendrites) and RAB7A 26 cells (from 3 independent cultures representing 93 dendrites).

Fig. 8 G-L

soma: (G-I) no Cre=122 cells; Cre-GFP=117 cells; Cre-GFP+WT RAB7A=124 cells; GFP+RAB7A-L8A=124 cells; Cre-GFP+Rab7A-F45A=122 cells.

--dendrites: (J-L) no Cre=53 cells; Cre-GFP=50 cells; Cre-GFP+WT RAB7A=49 cells, Cre-GFP+RAB7A-L8A=46 cells; Cre-GFP+Rab7A-F45A=46 cells. 4 independent cultures.

## Supporting information

manuscript

## Acknowledgments

This work was funded by NIH R01NS083378 to BW. We thank Dr. Paul Barnes for generously providing RAB7A^fl/fl^ mice (with permission from Dr. Aimee Edinger who initially made these mice).

## Notes

### Competing Interest Statement

The authors have declared no competing interest.

### Summary of Updates

We added live imaging track number data that we reanalyzed from published data in dynein inhibition experiments.

