## Supplementary material for "Late endosome transport by RILP-RAB7A promotes dendrite arborization": manuscript

**Supplementary Figures**

**SFig 1:** *Characterization of RILP-binding mutants RAB7A-L8A*

(A,B) Co-immunoprecipitation of Myc-RILP with GFP-RAB7A and its mutants from transfected HEK293 cells. HEK293 cells were co-transfected with myc-RILP and one of the following plasmids: GFP, GFP-RAB7A WT, GFP-RAB7A-T22N, GFP-RAB7A-L8A or GFP-RAB7A-F45A. 10% of the lysates were blotted to determine input levels for myc-RILP (anti-myc blot) or GFP-tagged constructs (anti-GFP blot). 90% of the lysates was used for a co-immunoprecipitation with GFP-Trap and blotted with anti-myc to detect the amount of bound myc-RILP. Only WT GFP-RAB7A co-immunoprecipitated myc-RILP. Three different experiments were carried out and quantified in (B).

(C-E) WT RAB7A (C), RAB7A-L8A (D), or RAB7A-F45A (E) were expressed in siRAB7 knockdown neurons together with myc-RILP. Both WT RAB7A and RAB7A-L8A are recruited to endosomes, but only WT RAB7A is able to recruit myc-RILP to endosomes. In RAB7A-L8A expressing neurons, myc-RILP remains cytosolic, consistent with loss of RILP binding by RAB7A-L8A. RAB7A-F45A is inefficiently recruited to endosomes and myc-RILP remains cytosolic. MAP2 (blue) indicates dendrites. All images are of DIV10 rat hippocampal neurons.

Yap et al. SFig.1

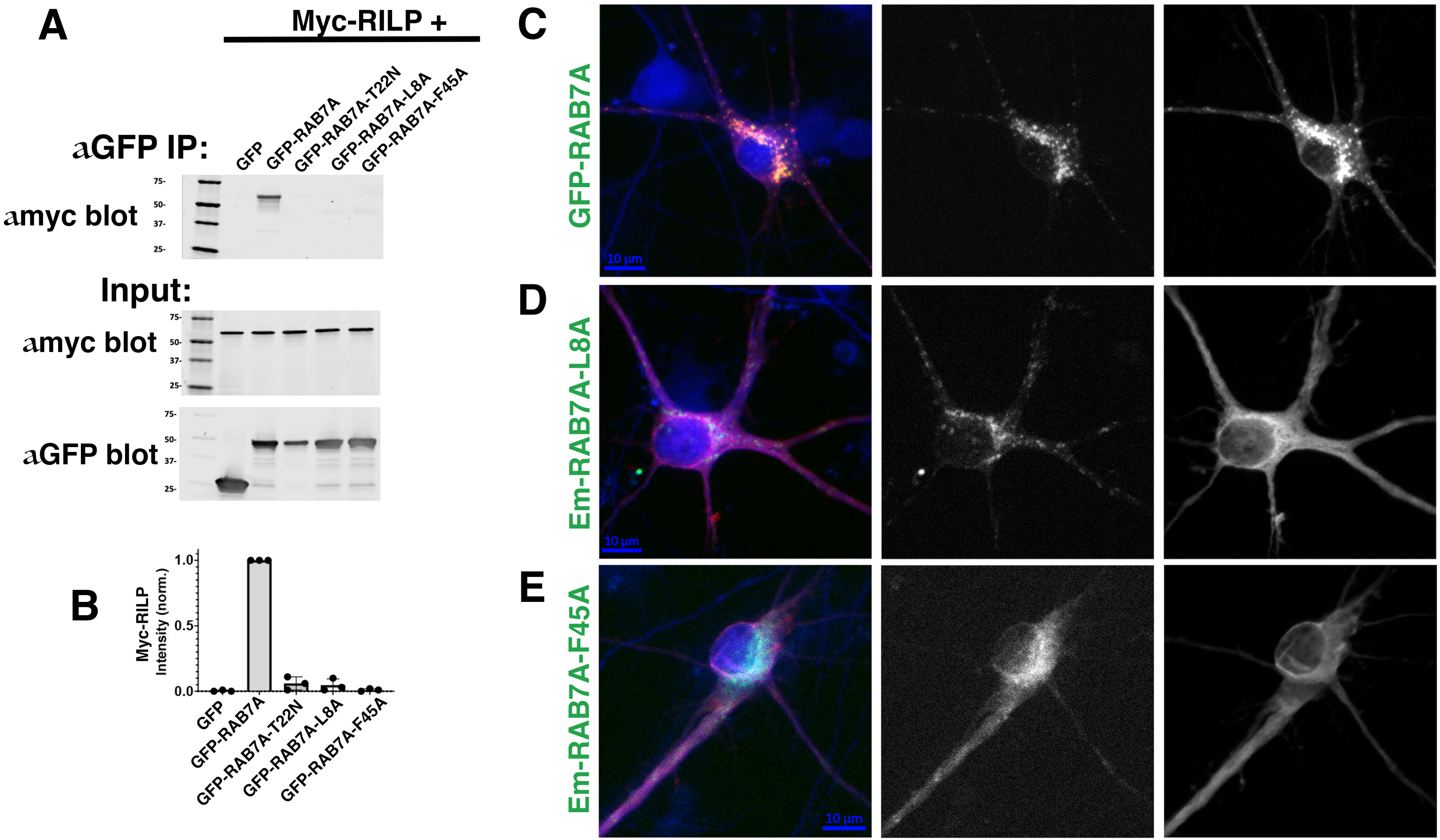

**SFig 2**: *Characterization of RILP-binding mutants RAB7A-F45A*

(A-B) Localization of Em-RAB7A-F45A (green) with endogenous markers of somatic endosomes (LAMP1: A; NSG2: B) and lysosomes (LAMP1: A; CatD: A; CatB: B) in DIV10 rat hippocampal neurons. MAP2 marks dendrites.

(C) Localization of Em-RAB7A-F45A (green) with endogenous markers of somatic early endosomes (Vps35 in red; EEA1 in blue) in DIV10 rat hippocampal neurons. For A-C, siRAB7 was co-transfected to minimize contribution to marker distribution from endogenous RAB7A.

(D) Em-RAB7A-F45A (green or cyan) localizes with FYCO1-FLAG (red) in some transfected neurons but is often largely cytosolic.

SFig.2

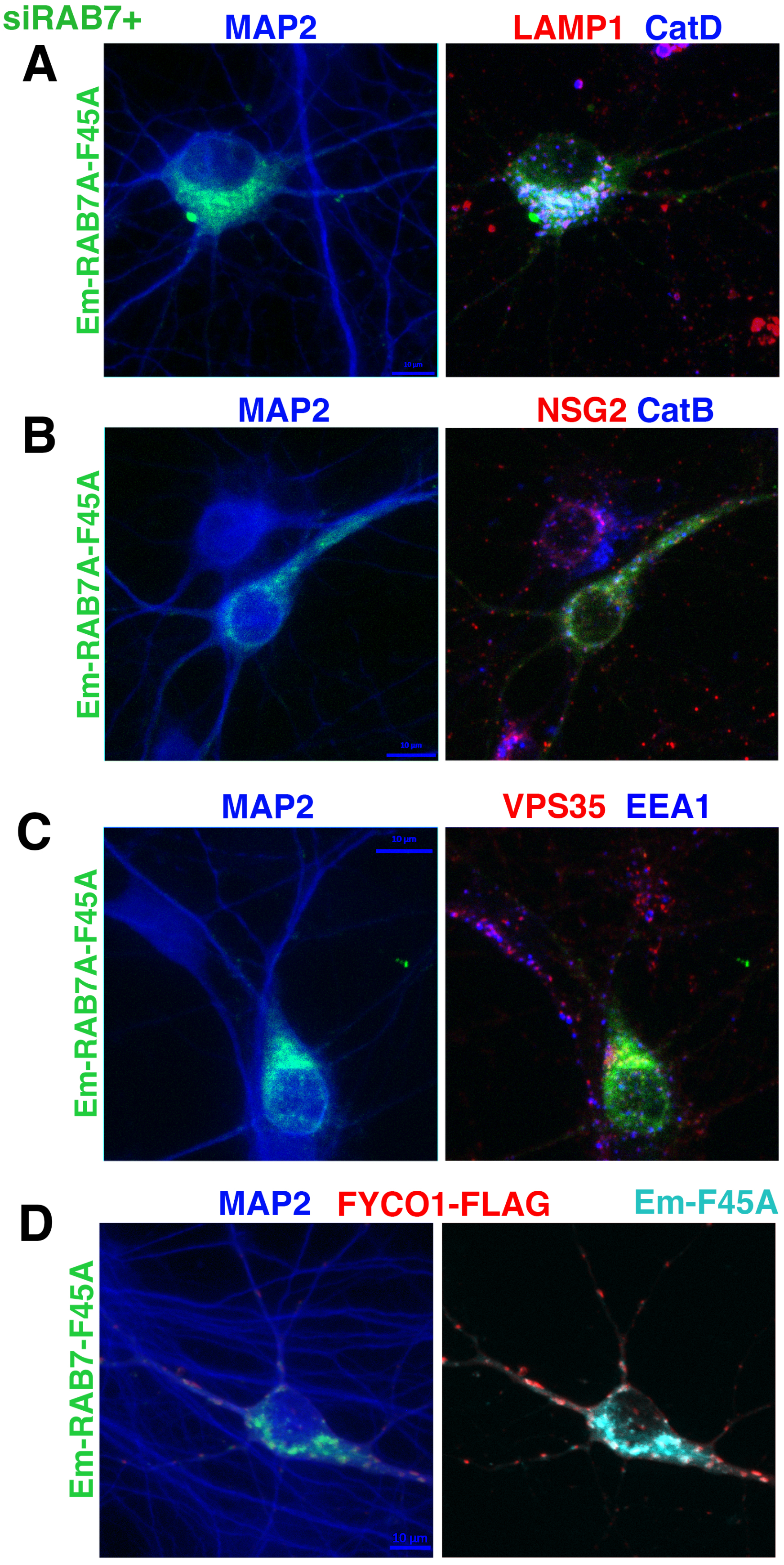
siRAB7+
A
MAP2
LAMP1 CatD
Em-RAB7A-F45A
B
MAP2
NSG2 CatB
Em-RAB7A-F45A
C
MAP2
VPS35 EEA1
Em-RAB7A-F45A
D
MAP2 FYCO1-FLAG
Em-F45A
Em-RAB7-F45A
10 µm

**SFig 3**: *Additional control data sets for dendrite analysis in Fig. 4*
(A-D) Additional comparison by Sholl analysis of DIV10 hippocampal neurons transfected with siCon or siRAB7 and different RAB7A constructs. Statistics: Multiple unpaired t-test. siCon-GFP 67 cell, siCon+WT RAB7A 73 cells, siRAB7+WT RAB7A 69 cells, siRAB7+DN-RAB7A 76 cells, siCon+DN-RAB7A 62 cells, siRAB7+GFP 73 cells, siRAB7+RAB7A-F45A 68 cells, siCon+RAB7A-L8A 73 cells, siCon+RAB7A-F45A 66 cells, siRAB7+RAB7A-L8A 72 cells.
(A) Sholl graphs of control conditions comparing siCon+GFP vs siCon+WT RAB7A vs siRAB7 + WT RAB7A rescue. The three conditions are not statistically different.
(B) Sholl graphs of RAB7A interference conditions comparing siRAB7+RAB7A-DN vs siCon+RAB7A-DN vs siRAB7+GFP. Expression of RAB7-DN in combination with either siCon or siRAB7 further diminishes dendrite complexity compared to siRAB7+GFP. * $p<0.05$. ** $p<0.01$.
(C) Sholl graphs addressing if RAB7mutants have a dendrite phenotype in the context of siCon. Both RAB7A-L8A and RAB7A-F45A have mildly reduced dendrite complexity in proximal branching. * $p<0.05$. ** $p<0.01$.
(D) Sholl analysis comparing rescue abilities of WT RAB7A vs RAB7A-DN vs RAB7A-L8A vs RAB7A-F45A. RAB7A-L8A and RAB7A-F45A are partial rescue in between RAB7A-DN and WT RAB7A. ** $p<0.01$. *** $p<0.001$. **** $p<0.0001$.

SFig. 3

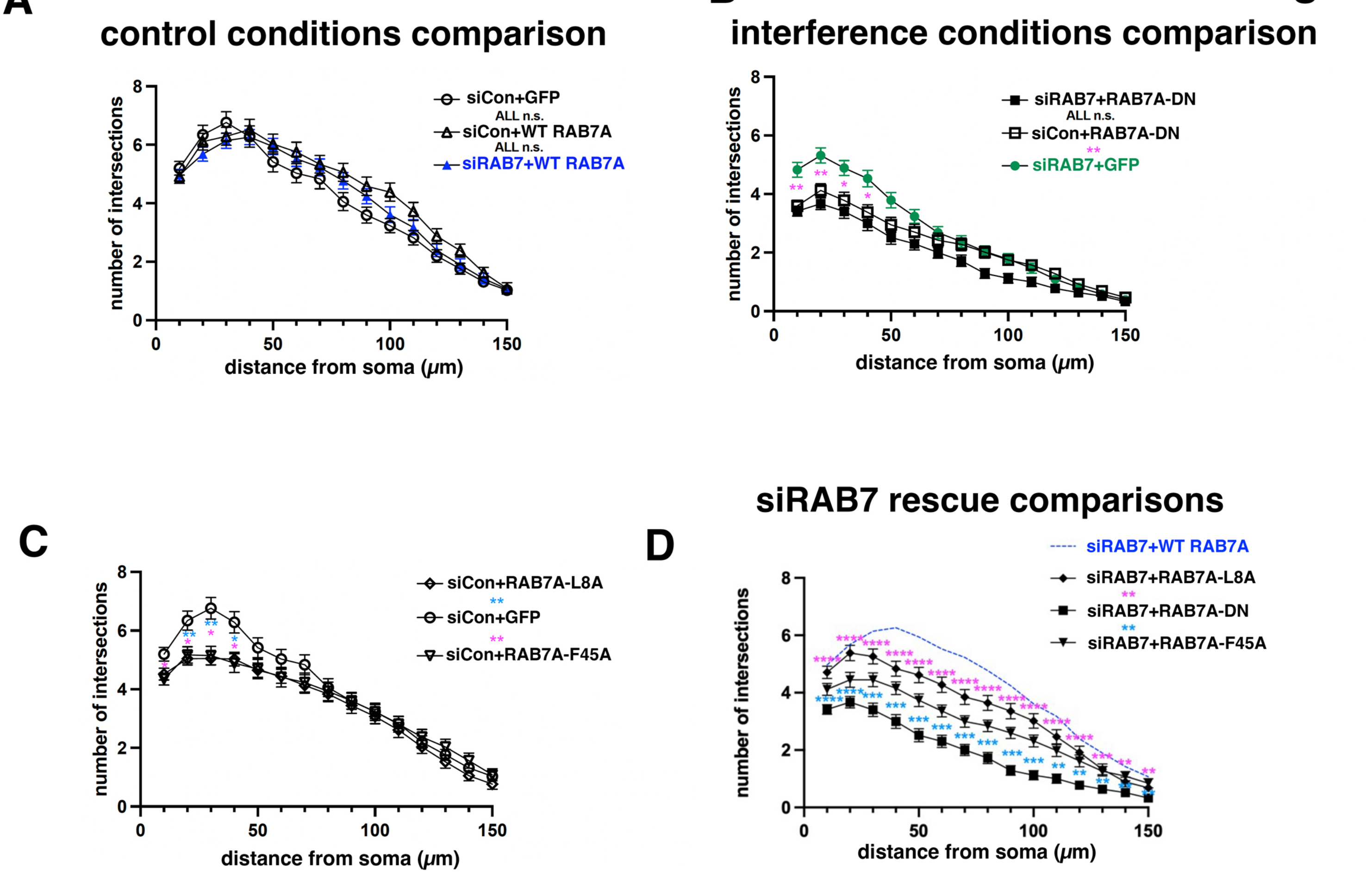

A
control conditions comparison
siCon+GFP
ALL n.s.
siCon+WT RAB7A
ALL n.s.
siRAB7+WT RAB7A
number of intersections
distance from soma (μm)
B
interference conditions comparison
siRAB7+RAB7A-DN
ALL n.s.
siCon+RAB7A-DN
**
siRAB7+GFP
number of intersections
distance from soma (μm)
C
siCon+RAB7A-L8A
**
siCon+GFP
**
siCon+RAB7A-F45A
number of intersections
distance from soma (μm)
siRAB7 rescue comparisons
D
siRAB7+WT RAB7A
siRAB7+RAB7A-L8A
**
siRAB7+RAB7A-DN
**
siRAB7+RAB7A-F45A
number of intersections
distance from soma (μm)
